# On specialists and generalists: niche range strategies across the tree of life

**DOI:** 10.1101/2022.07.21.500953

**Authors:** F. A. Bastiaan von Meijenfeldt, Paulien Hogeweg, Bas E. Dutilh

## Abstract

Generalists can survive in many environments whereas specialists have a limited distribution. Although a classical concept in ecology, niche breadth has remained challenging to quantify because it depends on an objective definition of the environment. Here, by defining the environment of a microbe as the community it resides in, we integrated information from over 22 thousand environmental sequencing samples to derive a quantitative social niche breadth score for all microbial taxa. At the level of genera, we explored niche range strategies across the tree of life. We found that generalists include opportunists that stochastically dominate local communities, while specialists are stable but low in abundance. Generalists have a more diverse and open pan genome than specialists, but we found no global correlation between niche breadth and genome size. Instead, we observed two distinct evolutionary strategies, where specialists have relatively small genomes in habitats with low local diversity, but relatively large genomes in habitats with high local diversity. Together, our global analysis shines a new, data-driven light on microbial niche range strategies.

---

Culture-independent sequencing studies have greatly expanded our understanding of the microbial world. They uprooted the tree of life ^1,2^, revolutionised our view of the human microbiome and virome ^3,4^, and advanced our comprehension of early evolution ^5,6^. By using standardised protocols across large numbers of samples ^7–10^, classical ecological questions can now be addressed on the global scale. A quintessential question is that of ecological niche breadth ^11^, the range of conditions in which an organism can live. Although the distinction between specialists and generalists is a fundamental property of life and its evolution, general mechanisms that determine niche breadth are poorly understood ^12^, and quantification has proven challenging ^13^.

Microbial niche breadth has been measured for specific aspects of the environment (e.g., temperature ^14,15^, pH ^16^, or nutrient dependence ^17,18^). General niche breadth definitions have been based on external habitat annotations. Some studies defined organisms that are present in many samples or predefined habitats as generalists, and rare organisms as specialists ^19–22^. Based on this definition, Sriswasdi et al. suggested an important evolutionary role for generalists in maintaining taxonomic diversity, with generalists having higher speciation rates and persistence advantages over specialists ^23^. Others defined the niche breadth of an organism by the uniformity of its distribution across habitats ^24^, suggesting that community assembly of specialists is driven by deterministic processes whereas for generalists neutral processes are more important ^25,26^. Notwithstanding these intriguing results, previous niche breadth studies have been sensitive to biases due to habitat definition and sample selection.

Microbiomes are sensitive biosensors capable of detecting geochemical gradients ^27^, disease state ^28,29^, or metabolites in a given niche ^30,31^. We thus reason that the vast collection of tens of thousands of environmental sequencing datasets that are available in the public domain ^32^ could be used to implement an unbiased, data-driven, and comprehensive niche breadth definition, based on community similarity between samples where microbial taxa occur. In this view, organisms that occur in compositionally similar samples are specialists, and organisms that occur in diverse samples are generalists. Using community similarity as a proxy for ecological range, we developed a social niche breadth (SNB) score that allowed us to quantify niche range for taxa at all taxonomic ranks and assess strategies for specialisation and niche range expansion across the prokaryotic tree of life.

## Results

### Social niche breadth captures global heterogeneity in microbial communities

To compare the niche breadth of microbial taxa, we devised and extensively benchmarked **(Supplementary results and discussion)** a social niche breadth (SNB) score that exploits the abundantly available meta’omics datasets derived from diverse environments around the world **(Fig. 1, Supp. table 1)**. First, we assessed the biome annotations of these datasets, as provided by the dataset submitters.

**Figure 1.**
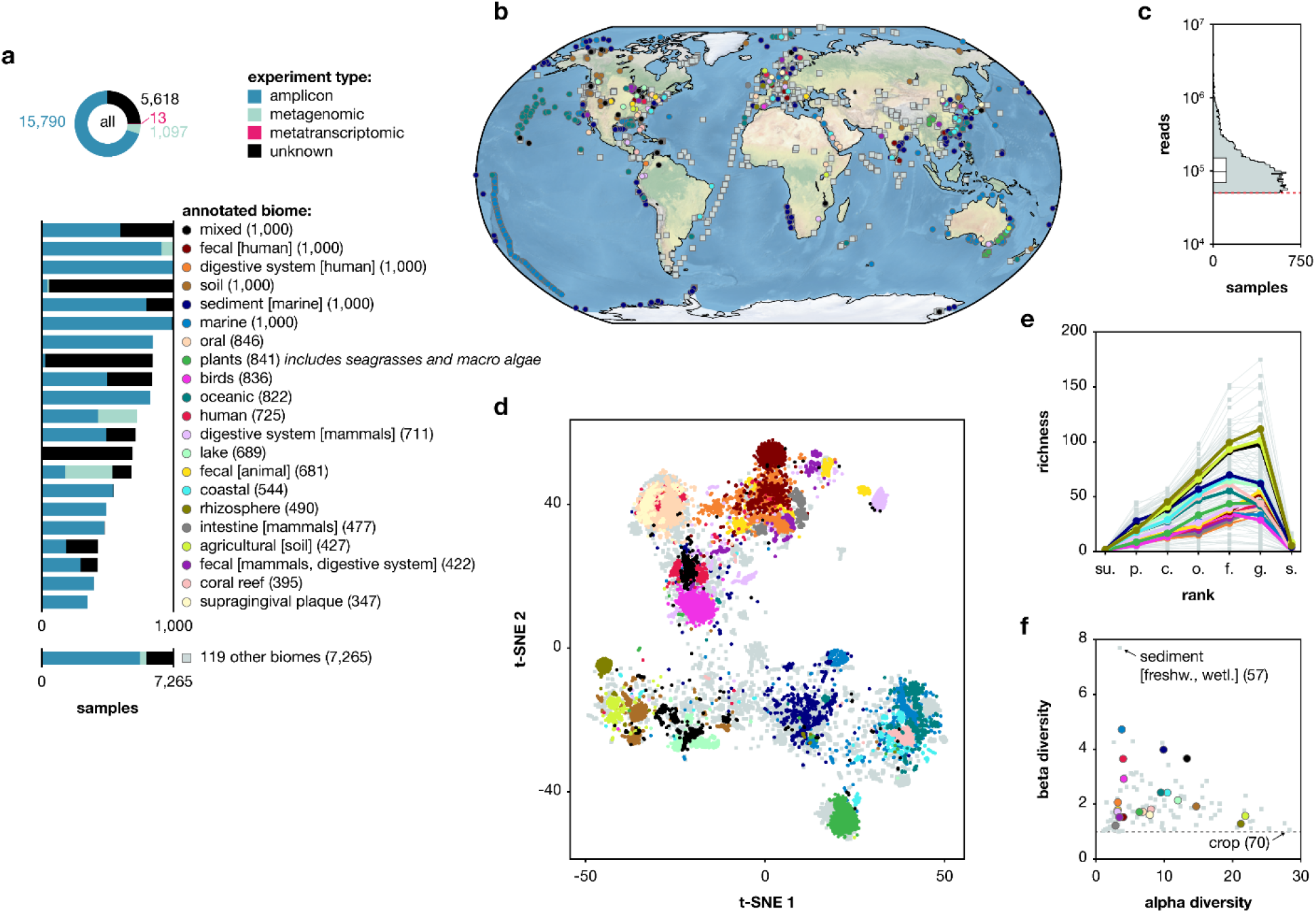
A diverse and global microbial dataset. (**a**) Samples are coming from vastly different annotated biomes and study designs. Numbers within brackets indicate the number of samples within the annotated biome. Annotated biomes with less than 347 samples are grouped as ‘other’ in this figure. For a hierarchical tree of all annotated biomes see **Supp. fig. 1b**. (**b**) Geographical distribution of samples. (**c**) Total number of taxonomically annotated reads per samples. The red line shows the cut-off of 50,000 reads that was one selection criterion. (**d**) Samples from similar annotated biomes cluster together based on taxonomic profile in a t-distributed stochastic neighbour embedding (t-SNE) visualisation (perplexity 500), with the same ecological dissimilarity measure used as for SNB, namely Spearman’s rank correlation coefficient (0.5 − (*ρ*/2)) of known taxa at rank order. Samples are separated by host-association and salinity. For a PCoA visualisation of the same data and positions of all 140 annotated biomes on the PCoA see **Supp. fig. 2** and **Supp. fig. 3**, respectively. Most samples from the plants biome are derived from seagrasses and macro algae from kelp forests, which is why they cluster near marine samples. (**e**) Taxa richness differs per annotated biome and taxonomic rank. The low number of annotated species is a consequence of a relatively unexplored biosphere. (**f**) Annotated biomes with high alpha diversity have low beta diversity.

These annotations highlighted the main drivers of microbiome composition **(Fig. 1d, Supplementary results and discussion)**, including salinity (t-SNE 1) and host-association (t-SNE 2) ^10,33–35^. The 22,518 samples covered a total of 140 annotated biomes that differed markedly in within-sample (alpha) and between-sample (beta) diversity. Notably, annotated biomes with high alpha diversity such as soils had low beta diversity **(Fig. 1f)**, implying a low turnover rate and stable community composition of highly diverse habitats. Most annotated biomes have low beta diversity reflecting consistent microbial composition. Nevertheless, annotated biome definitions are arbitrarily delineated and may be subject to human error. For example, the plants biome includes both freshwater plants ^36–38^ and seagrasses ^39^, as well as macro algae from kelp forests ^40^ **(Supp. table 1)**. Also, it remains difficult to quantify the degree of similarity between categorical biomes. We used the observation that microbiomes are biosensors and developed SNB that captures the compositional heterogeneity of the samples where a taxon is found to quantify niche breadth. This approach accounts for database biases, as some environments are more frequently sampled than others **(Fig. 1a, Supplementary results and discussion)**. Indeed, taxa that are detected in the same number of samples may have very different SNB **(Fig. 2a-c)**.

**Figure 2.**
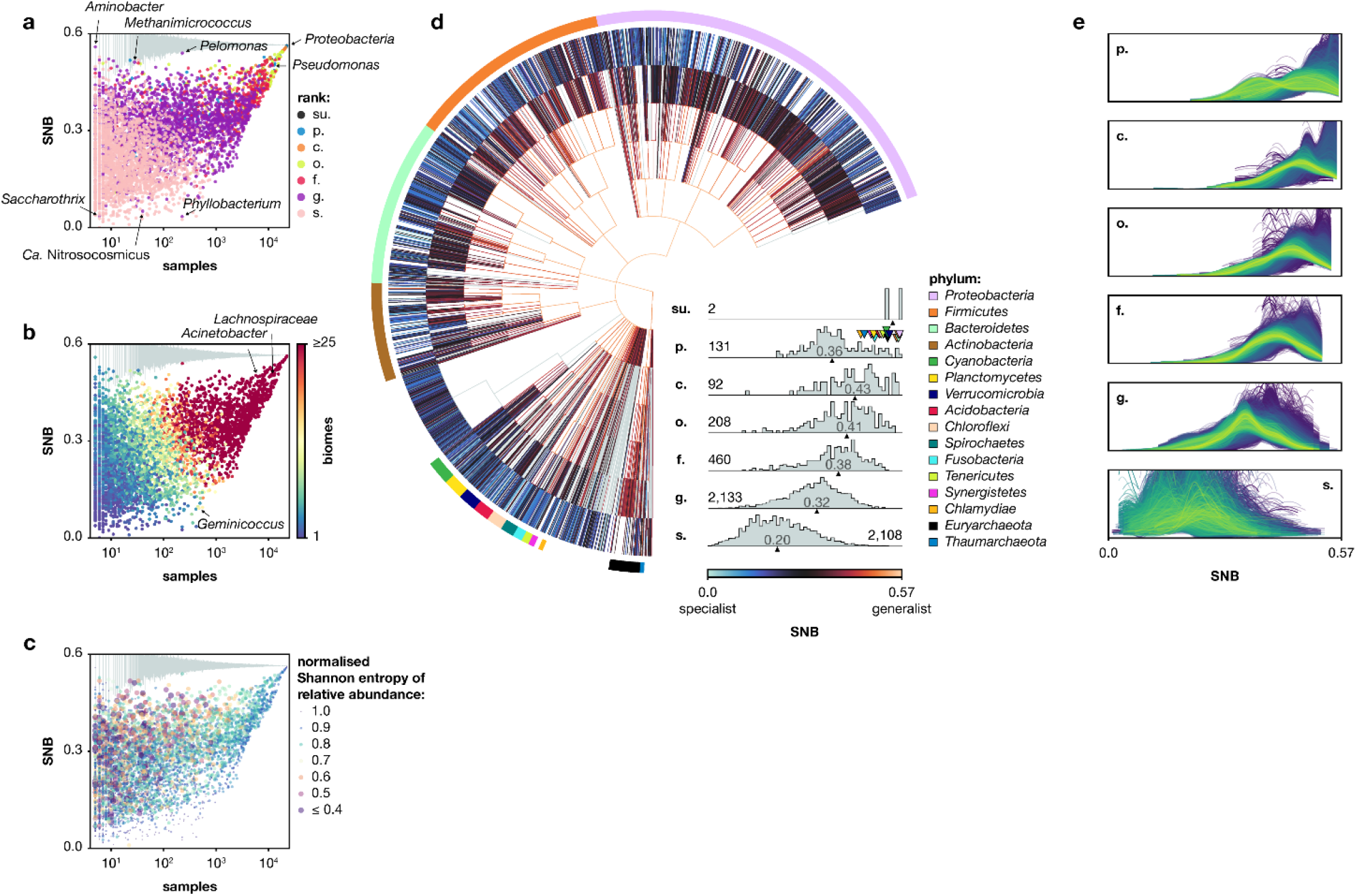
Social niche breadth across the tree of life and across samples. (**a**,**b**,**c**) Relation between SNB and the number of samples in which a taxon is found. Grey bars on top show the range of social niche breadth of imaginary taxa that are present in 100 randomly picked subsets of samples of the specific size. The locations of some outlier taxa are indicated. Normalised Shannon entropy of relative abundances across samples is used as shading for each taxon in panel c. Both colour-coding and size of markers represents the Shannon entropy. Note that higher entropy is indicated with smaller markers. Relative abundance across samples is more constant for specialists than for generalists. (**d**) SNB of taxa throughout the tree of life. Grey colour-coding of branches represents unnamed taxa at that rank. Distribution of SNB at different taxonomic ranks is shown as histograms, numbers on the distributions show the number of taxa at that rank. Upward pointing triangles below and numbers within distributions indicate the median value for that rank. Downward pointing triangles above the phylum distribution indicate the value of the colour-coded phyla. (**e**) Distribution of SNB within samples at different taxonomic ranks. Blue lines indicate samples with low alpha diversity and yellow lines indicate samples with high alpha diversity.

### Niche breadth across the tree of life and at different taxonomic ranks

To investigate the distribution of social generalists and specialists throughout the prokaryotic tree of life, we calculated SNB for taxa at all ranks **(Fig. 2d, Supp. table 2)**. For the vast majority of taxa, the SNB score is lower than expected based on random permutations **(Fig. 2a-c)**, indicating that all microbes are specialists to some extent because they occur in a non-random subset of all samples. As expected, exceptions to this rule include the high-ranking superkingdom *Bacteria* and phylum *Proteobacteria* which are widespread, occurring in 22,295 and 22,211 of the 22,518 samples, respectively. Notably, while there is a clear positive correlation between SNB and the number of samples where a taxon occurs **(Fig. 2a-c)**, very rare taxa such as *Aminobacter* (5 samples) and *Methanimicrococcus* (28 samples) still have a high SNB (SNB = 0.56 and SNB = 0.51, respectively). Alternatively, some taxa that are found in many samples have a relatively low SNB because these samples are very similar in composition, for example *Phyllobacterium* (226 samples, SNB = 0.03) and *Geminicoccus* (473 samples, SNB = 0.09).

The distribution of SNB differs per taxonomic rank, where high-ranking taxa tend to be generalists and low-ranking taxa specialists **(Fig. 2d,e)**. High-ranking taxa can be generalists because they contain subtaxa that are specialists in different communities, or because the subtaxa are also generalists. For example, genera of the generalist family *Flavobacteriaceae* and species of the generalist genus *Prevotella* are relatively specialised for their respective rank. *Flavobacteriaceae* has an SNB of 0.52, and its genera a median SNB of 0.28, and *Prevotella* has an SNB of 0.37, and its species a median SNB of 0.14 (see **Fig. 2d** for the median SNB of all taxa per rank). Genera within the generalist family *Lactobacillaceae* (SNB = 0.46) on the other hand have a median SNB of 0.45. In addition, generalist taxa often have more subtaxa than specialist taxa **(Supp. fig. 14)**. This suggests that the diversity of taxa, as currently represented by taxonomy, reflects their ecological range well. Indeed, the four best-studied phyla, *Proteobacteria, Firmicutes, Bacteroidetes*, and *Actinobacteria*, that together cover 97% of cultured prokaryotic species ^41^, have a higher SNB than others **(Fig. 2d)**.

Candidate phyla have a low SNB compared to established phyla **(Fig. 2d, Supp. fig. 15)**. Because the candidate phyla contain less described subtaxa than established phyla, the distribution of SNB on phylum rank is more skewed towards specialism than class, order, and family ranks **(Fig. 2d)**. Only recently discovered, candidate phyla generally require specific growth conditions and are thus difficult to cultivate, consistent with their low SNB. Several candidate phyla including the bacterial Candidate Phyla Radiation and DPANN archaea may consist of obligate symbionts of specific hosts ^2^. Whereas it was recently shown that consortia of obligate symbionts can grow on a wider range of carbon sources than their individual members and thus expand their metabolic niche ^42^, the individual microbes in these consortia are specialists from the SNB perspective as they require specific partners in their local communities.

Taxa with high and low SNB are dispersed throughout the tree of life, indicating that specialisation and niche range expansion happened independently numerous times in evolution. Phyla with relatively specialised genera include *Proteobacteria, Bacteroidetes, Actinobacteria, Cyanobacteria, Planctomycetes, Acidobacteria*, and *Chloroflexi*, whereas *Firmicutes, Tenericutes*, and *Euryarchaeota* have relatively generalised genera **(Supp. table 3)**. Taxa with relatively low SNB for their ranks include known specialists such as *Christensenella* ^23^ (SNB = 0.24), but also the taxa *Pelagibacteraceae* (SNB = 0.25) and *Prochlorococcus* (SNB = 0.22), which hold some of the most abundant organisms on Earth ^43,44^. These taxa, known for their highly streamlined genomes ^45^, are found in aquatic samples with a uniform microbial composition **(Supp. fig. 5b)** and thus have a low SNB. The genus *Roseobacter* (SNB = 0.30), whose members are considered marine generalists with large genomes and a versatile metabolism ^46,47^, is found in more diverse samples **(Supp. fig. 5b)** and has an SNB close to the median of all genera. At the generalist end of the spectrum are ubiquitous taxa like *Acinetobacter* (SNB = 0.50) and *Pseudomonas* (SNB = 0.50) **(Supp. fig. 5b)** (but see ^48^). *Lachnospiraceae* (SNB = 0.50) and its genera **(Supp. table 3)**, obligate anaerobes that were previously regarded as specialists ^23^ also have a high SNB, highlighting the diversity of their anaerobic habitats.

### Generalists compete for dominance, specialists are stable at low abundance

Next, we set out to find patterns in social niche breadth. We focussed our analysis on genera because they balance a high taxonomic resolution with a good representation in the dataset **(Fig. 1e)** and show a broad range in SNB values **(Fig. 2d)**, allowing for a comprehensive investigation of niche range strategies.

It has been suggested that generalists, being jack of all trades, can be master of none, while specialists are adapted to become dominant within their habitats ^49^. Niche range may thus reflect a trade-off, where specialists gain local dominance at the expense of ecological versatility. Alternatively, computational models of microbial metabolism have suggested that metabolically flexible generalists have faster growth rates than specialists ^50^. We correlated SNB with local abundance and found that generalists are dominant in most annotated biomes, exceptions including marine organisms like corals, seagrasses, and macro algae **(Fig. 3a)**. Thus, generalist genera often outcompete specialists in their local communities, disputing the expected trade-off mentioned above. Local dominance of generalists has previously been observed in specific environmental settings like highly dynamic sandy ecosystems ^51^. Some soil microbes are both abundant and ubiquitous ^52^, and only ∼500 dominant phylotypes (i.e. 2%) represent >40% of all soil bacteria ^53^. Our results show that these observations reflect a general pattern wherein generalists are dominant.

**Figure 3.**
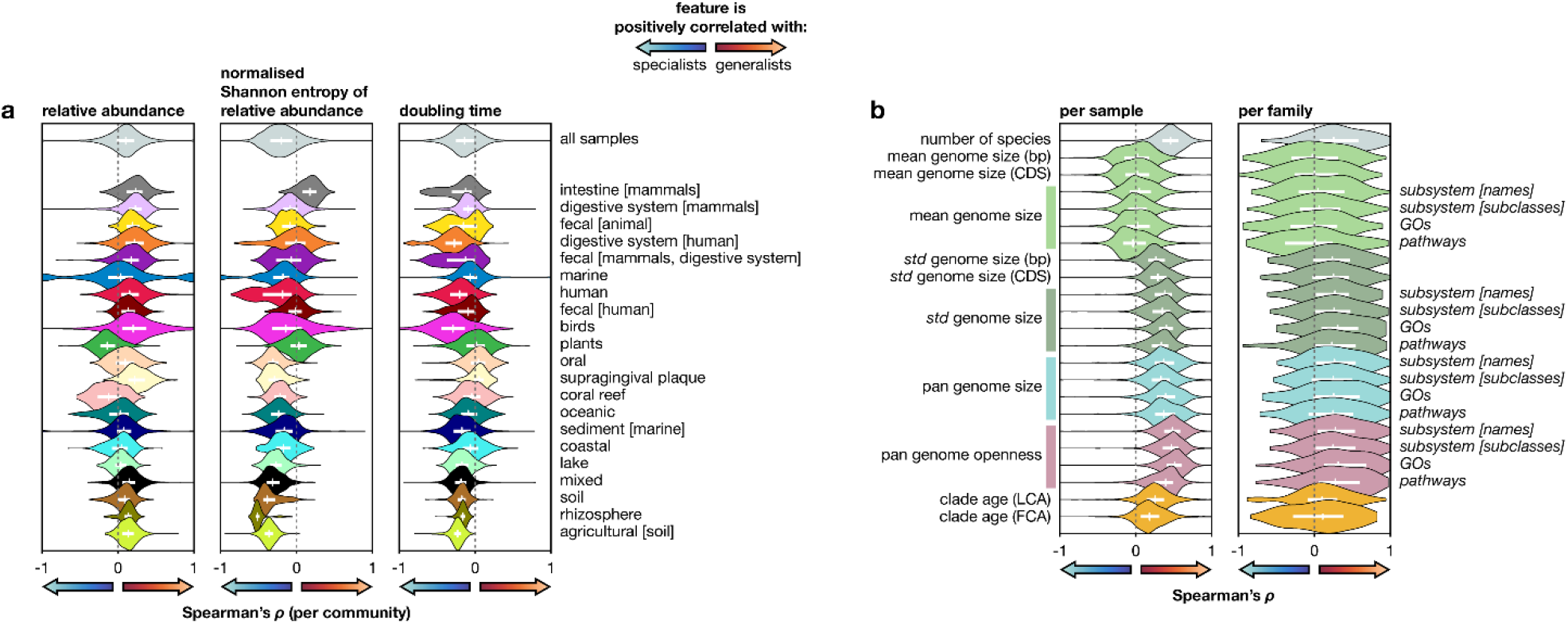
Ecological and genomic features correlated with SNB. (**a**) Spearman’s rank correlation coefficient (*ρ*) per sample between SNB and features related to local dominance on the rank genus. Violins depict the distribution of *ρ* across all samples or those from the annotated biomes with the most samples. Annotated biomes are arranged according to alpha diversity. Positive values indicate that the feature is positively correlated with generalists, and negative values indicate that the feature is positively correlated with specialists. Doubling time estimates are from the EGGO database. (**b**) *ρ* per sample and per family between SNB and genomic features on the rank genus. Violins depict the distribution of *ρ* across all samples, and across all families with at least 5 genera. Genome sizes are given in base pairs (bp) and number of coding regions (CDS). Genomic measures with annotations to the right are in number of unique functions for that specific functional universe. Genome size estimates for a genus are based on the genome size of its species, which is defined as the mean size of all strains for bp and CDS measures, and as the majority set of functions of all strains for the functional universe measures. Pan genome openness is total pan genome size divided by mean genome size. Correlations with time to the Last Common Ancestor (LCA) and First Common Ancestor (FCA) are based on the TimeTree database. Lines within violins show interquartile range and median.

Whereas local communities are typically dominated by generalists, the relative abundance of generalists is more variable across samples than that of specialists, whose abundance is relatively stable. This is evident when comparing the niche range of organisms that locally co-occur within samples, where generalists have a higher variability of relative abundance than specialists **(Fig. 3a)**. It is also evident for taxa across the tree of life, where specialists have an even relative abundance while taxa with a high variability of relative abundance are generalists **(Fig. 2c)**. Social niche breadth may thus partially reflect the classical distinction between *r*-strategists and *K*-strategists. Specialists have a low but constant abundance near carrying capacity (*K*-selected), and some (but not all) generalists are opportunistic taxa that reach high relative abundance when circumstance permits (*r*-selected). To test this hypothesis, we compared SNB of microbes to their predicted maximal growth rates ^54^ **(Fig. 3a)**, and confirmed that, within samples, generalists have shorter doubling times than specialists. These results support the idea that generalist genera include more opportunistic growers than specialist genera.

### Niche breadth reflects genomic heterogeneity

Next, we used our dataset to assess the suggestion that generalist have large genomes that encode many functions, reflecting a versatile metabolism that allows them to colonise diverse habitats ^21,23^. For example, bacteria that are found in a diverse range of habitats encode more extracellular proteins than bacteria that are restricted to few habitats ^22^, and habitats with temporal variation may select for larger genomes ^55^. In contrast, specialisation may be associated with a reduction in genome size due to loss of unnecessary genes, as has been observed in members of the phylum *Planctomycetes* transitioning from soil to freshwater habitats ^56^, or genome streamlining ^57^, which is common in oligotrophic marine waters ^58,59^. Genomic versatility of high-ranking taxa, reflected in a large pan genome ^60,61^, may either result from small yet diverse genomes in individual subtaxa (open pan genome), or from genomically versatile yet functionally similar strains (closed pan genome). We set out to identify genomic features associated with social niche breadth using publicly accessible genome sequences ^62^ **(Supp. table 4)**, by correlating genomic features of genera with their SNB. We compared genera within samples **(Fig. 2e)** for an ecological view, and within their taxonomic families for an evolutionary view. Both perspectives gave qualitatively similar results (**Fig. 3b**; see below), indicating that genomic signatures of SNB are generalisable across habitat and phylogeny. Although the number of samples is larger than the number of families, the correlation between genomic features and SNB is more consistent within samples than within families, possibly suggesting that ecology is a stronger driver of (pan) genome evolution than phylogenetic history ^63^.

When comparing taxa across all samples, we found no consistent correlation between SNB and genome size, whether measured in number of nucleotides, genes, or unique functions **(Fig. 3b)**. We do, however, observe that the genomes in generalist genera are more variable in size than the genomes in specialised genera, as seen in their standard deviation. Moreover, the pan genomes of generalist genera contain more functions, in line with theoretical models that suggest that the ability to migrate to new niches is associated to pan genome size ^64^. The pan genome size of microbes may be positively associated to effective population size ^65^, which may be larger for generalists. The same study found that rapidly growing microbes have large effective population sizes, in line with our earlier discussed observation that opportunistic growers are generalists. Finally, the pan genomes of generalists are more open than those of specialists. Notably, these results did not depend on the higher number of species in generalist genera **(Supp. fig. 16)**.

In conclusion, species in specialist genera are genomically more similar than species in generalist genera. The correspondence between community heterogeneity and genomic heterogeneity confirms the strong association between ecological and genomic diversification. To further explore this diversification we correlated SNB with clade age ^66^. It was previously suggested that generalist species are evolutionary younger than specialist species ^23^. Our data do not allow analysing these trends at the species rank, but at the genus rank we found that specialists are younger than generalists **(Fig. 3b)**. Together, these results support a model of continuous diversification, where old generalist clades that share a diverse pan genome may invade new niches, leading to the emergence of specialised subtaxa.

### Two contrasting genomic niche range strategies

As discussed above, SNB was not consistently associated with genome size. However, we did observe a habitat-dependent relation between genome size and niche range **(Fig. 4a)**. We found two contrasting strategies that broadly depended on the alpha diversity of the samples **(Fig. 4b)**. In samples with low diversity, including most animal-associated and saline habitats, there is a positive correlation between genome size and SNB. In contrast, in high diversity samples, that include most free-living non-saline habitats and the rhizosphere, the correlation is negative. Because databases contain a majority of samples from animal-associated and marine habitats with relatively low diversity **(Fig. 4c)**, and because genome size estimates are often based on cultivated microbes that differ markedly from environmentally derived genomes ^67^, previous suggestions of a positive correlation between genome size and niche range are likely biased.

**Figure 4.**
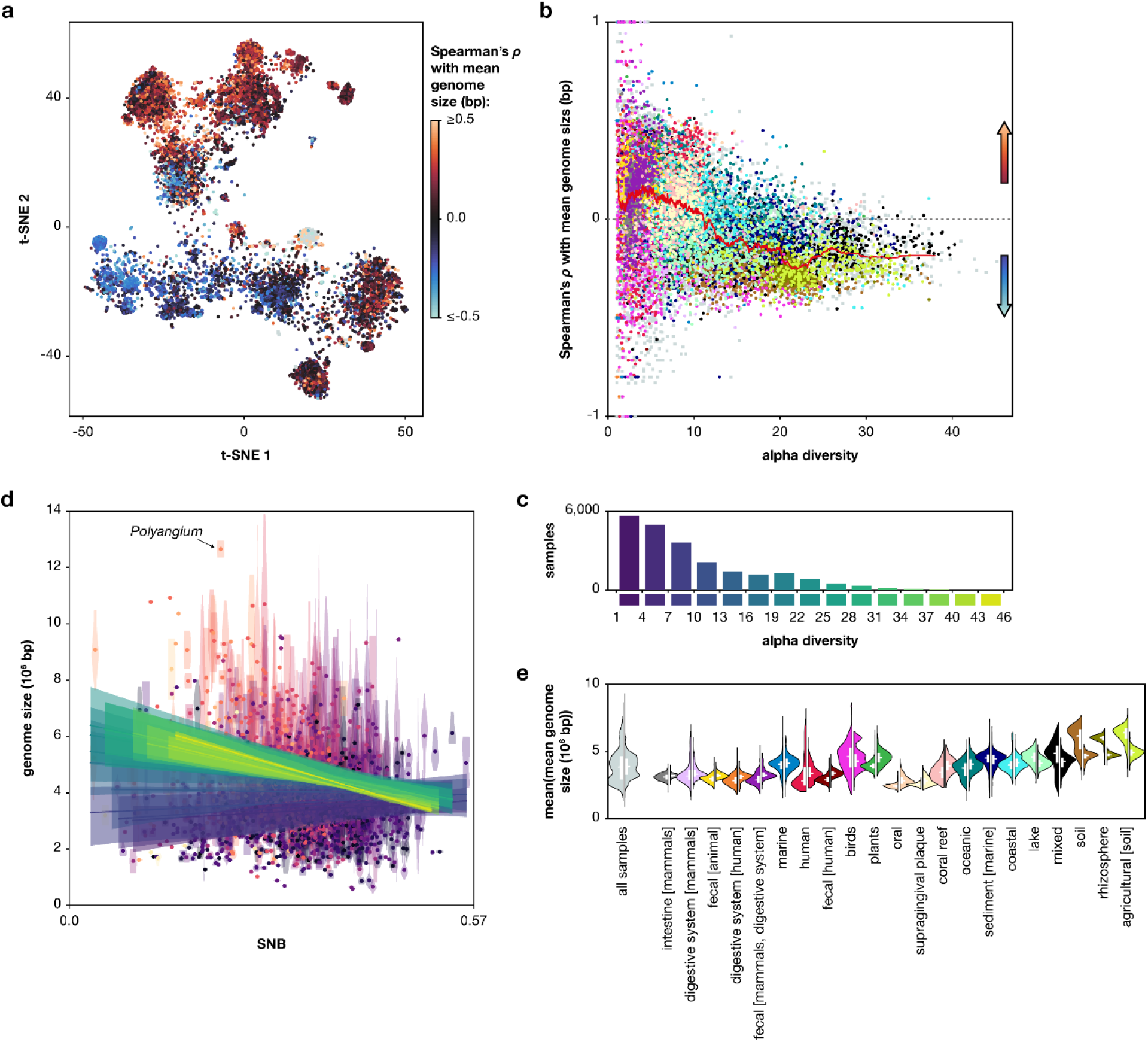
Contrasting genomic niche range strategies. (**a**) Spearman’s rank correlation coefficient (*ρ*) per sample between SNB and mean genome size on the rank genus plotted on the t-SNE **(Fig. 1d)**. Positive values indicate association with generalists, and negative values indicate association with specialists. (**b**) *ρ* as a function of the alpha diversity of the sample. Colour-coding represents annotated biomes, see panel e. The red line is the moving median with a window size of 250 samples. (**c**) Number of samples per bin of alpha diversity. Bins have the same colour as in panel d. (**d**) SNB versus genome size in base pairs on the rank genus. Violins show the distribution of genome sizes of species within a genus, dots are the mean. Colours of the violins and dots show the alpha diversity of the samples in which the genus is found, with lighter colours representing higher diversity than darker colours. Lines depict the mean of linear regression lines of all samples in a specific bin of alpha diversity, see panel c. Shaded areas show the interquartile range of the regression lines. (**e**) Violins depict the distribution of mean genome size of the top 25% specialist (left) and top 25% generalist (right) taxa per sample across all samples or those from the annotated biomes with the most samples. Annotated biomes are arranged according to alpha diversity. Lines within violins show interquartile range and median.

In low diversity habitats, generalists have large genomes that may encode the functions needed to utilise many different resources, while specialists (‘low diversity specialists’) have the smallest genomes known **(Fig. 4d,e)**. Coding density, a signature of genomic streamlining ^57^, is significantly higher in generalists than in low diversity specialists (p < 0.001, one-tailed T-test, measured in number of coding sequences per base pair). This suggests that genome streamlining is not the common route to genome reduction in low diversity specialists. Instead, their small genomes could reflect specialisation to habitat-specific metabolites and the loss of genes through drift ^56^. In addition, cooperating specialists could supplement each other’s nutrient requirements ^68^, or they could depend on the co-occurring generalists.

Although the genomes of generalists are large compared to those of low diversity specialists **(Fig. 4d)**, the genomes in low diversity samples are still moderate in size compared to the large genomes in high diversity samples **(Fig. 4e)**. Strikingly, we see the opposite correlation between genome size and niche range in high diversity samples, where specialists (‘high diversity specialists’) have larger genomes than co-occurring generalists **(Fig. 4b)**. The largest known prokaryotic genomes belong to high diversity specialists **(Fig. 4d)**, for example the genus *Polyangium* with a mean genome size of 12.7Mb. Selection may favour large genomes in habitats with diverse but scarce nutrient availability where slow growth is no disadvantage like in soils ^69,70^. Moreover, in contrast to the earlier mentioned cooperative taxa, microbes in competitive consortia carry many metabolic functions ^68^. High diversity specialists may thus reflect a competitive metabolism. Generalists in these habitats may use metabolites that are irregularly available and rapidly depleted, consistent with their opportunistic nature and variable occurrence. Alternatively, they could exploit metabolic byproducts generated by the specialists ^18^. Regardless of the mechanism, it appears that adaptation of high-diversity specialists to their habitats by genome expansion decreases their competitiveness in differing communities.

### Generalist pan genomes reflect fluctuating habitats, low and high diversity specialists differ in metabolic adaptations

To further characterise generalist and specialist microbes, we explored differences in genomic content by dividing all genera into two groups based on SNB **(Fig. 5a)** and performing Gene Set Enrichment Analysis (GSEA) ^71^ on the genus-level pan genomes. We performed two GSEAs, one with all genera comparing specialists to generalists **(Fig. 5b, Supp. table 5)**, and one comparing low diversity specialists to high diversity specialists **(Fig. 5c, Supp. table 6)**.

**Figure 5.**
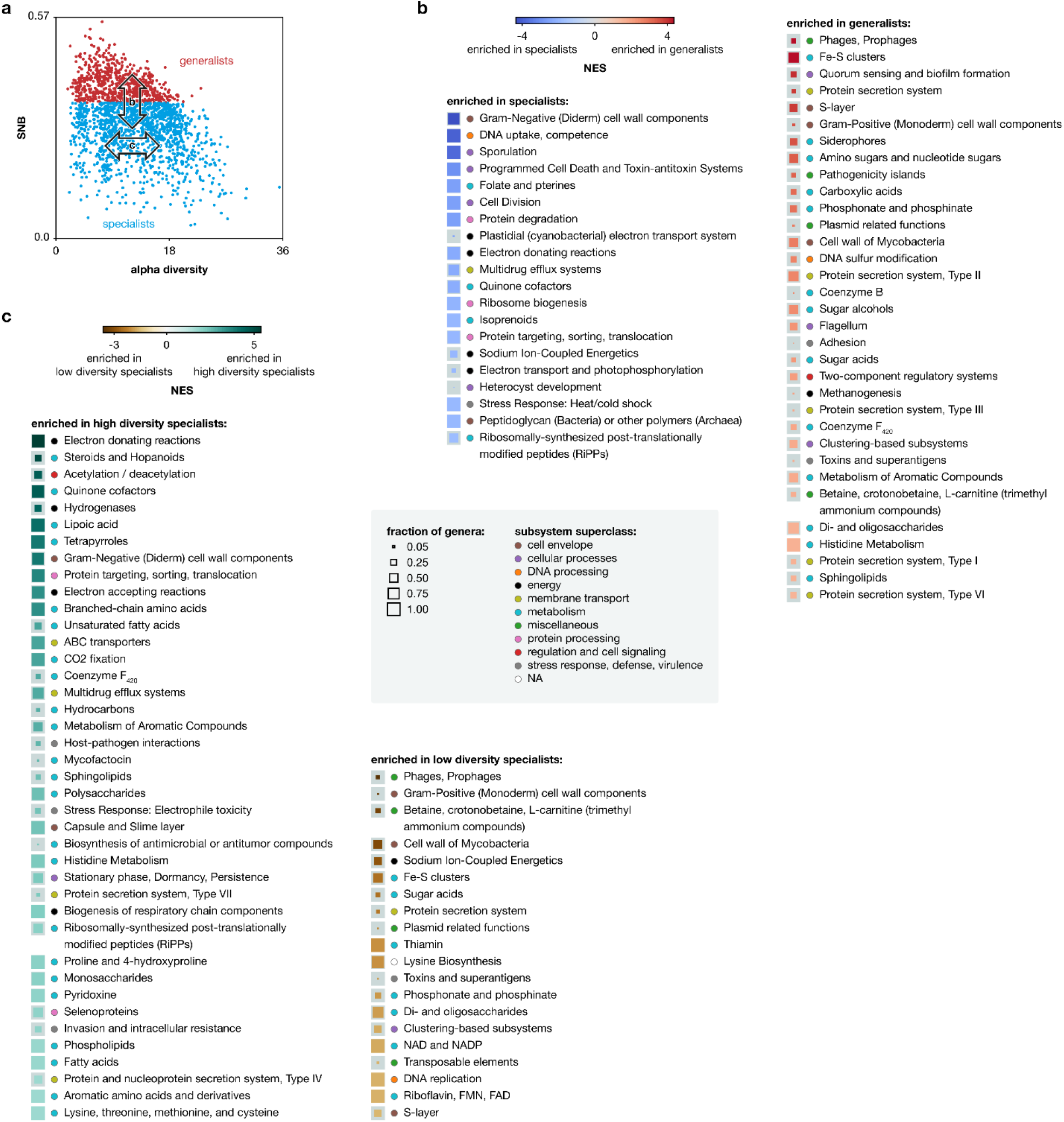
Functional characterisation of generalists and low and high diversity specialists. (**a**) Division of genera in specialists (SNB < 0.35) and generalists (other). (**b**) GSEA on the pan genome of all genera sorted by SNB, on the level of subsystem subclasses. (**c**) GSEA on the pan genome of specialist genera sorted by alpha diversity, on the level of subsystem subclasses. Enriched functions (FDR < 0.1) in panels a and b are sorted according to normalised enrichment score (NES), size of the square markers indicates the fraction of genera in that analysis having the function. The full GSEA information, including analyses of other functional universes (subsystem names, GOs, and pathways) can be found in **Supp. table 5** and **Supp. table 6**.

Fewer genomic functions were enriched in specialists than in generalists (false discovery rate (FDR) < 0.1, **Fig. 5b**). Specialist-enriched functions include energy related processes, and some specifically related to *Cyanobacteria*, such as heterocyte formation which is involved in nitrogen fixation in one evolutionary lineage of the phylum ^72^. Most enriched functions were present widely in pan genomes of both generalist and specialist genera **(Fig. 5b)**, suggesting that the smaller pan genome size of specialists does not involve consistent loss of functions. The absence of specific enriched specialist functions highlights that there is not a single type of specialist, but instead different specialists each have their own specific functions. In contrast, functions enriched in generalists **(Fig. 5b)** are more pronounced and reflect an investment in proteins related to species-species interactions, observation, and response to a fluctuating environment. Out of the 33 generalist-enriched functions, 13 were related to metabolism, including functions associated with secondary metabolites such as Coenzyme F_420_^21,73^. Other functions include quorum sensing and biofilm formation, adhesion, locomotion via the flagellum, and functions concerning the cell envelope and transport across it (S-layer, protein secretion systems, siderophores). Pathogenicity islands could point to opportunistic interactions with eukaryotic host organisms. Finally, functions associated with genome fluidity such as pro(phages) and plasmid-related functions are also enriched in generalists, highlighting the mechanisms by which they keep an open pan genome, allowing them to colonise diverse niches.

Comparing low diversity specialists to high diversity specialists **(Fig. 5c)**, we observed several specific metabolic adaptations to these different types of habitats. Half of the 60 enriched functions in the GSEA were related to metabolism. High diversity specialists have more enriched functions than low diversity specialists. For example, functions associated with stationary phase, dormancy, and persistence are enriched in high diversity specialists, consistent with slow growth and persistence in soil. Moreover, functions related to lipid metabolism (e.g. steroids and hopanoids, (unsaturated) fatty acids, sphingolipids, and phosopholipids) are enriched in high diversity specialists. Low diversity specialists, like generalists, also contain some functions associated with genome fluidity (transposable elements, (pro)phages, and plasmid related functions), suggesting that their genomes, although small in size, may still be in flux.

## Discussion

We present a social niche breadth score for microbial taxa that is based on the community similarity of the samples where they occur. Integrating information from over 22 thousand samples, SNB represents a global and comprehensive view on niche range across the microbial tree of life. With continued and ever deeper sequencing efforts and associated expansion of public databases, the environmental and taxonomic resolution of our picture of the microbial world increases, as does our understanding of the processes shaping microbial niche breadth. Contrary to earlier suggestions, we found that most habitats are dominated by generalists. Specialists occur at low but stable abundances.

Generalist genera are older than specialist genera and depend on large and open pan genomes that allow them to adapt to different habitats. Individual genome size and SNB are differentially related depending on the diversity of the habitat, generalists having larger genomes than specialists in low diversity habitats, and smaller genomes than specialists in high diversity habitats. High diversity specialists may need a large genetic repertoire as they are continually exposed to many different interaction partners and possibly high environmental variability at small spatial scales. Low diversity specialists have decreased genome sizes due to loss of unnecessary functions. Large genomes may thus reflect increased environmental versatility in two different settings. In low diversity habitats, generalists are relatively versatile, as they can survive in a range of different communities. In high diversity habitats, specialists are relatively versatile, allowing them to persist in their local complex community. Since generalists and specialists are dispersed throughout the tree of life, these genomic adaptations have repeatedly occurred and represent fundamental eco-evolutionary processes.

## Methods

### Sample selection

We downloaded taxonomic profiles deposited in the MGnify microbiome resource ^32^ on 20 August, 2019. MGnify contains taxonomic profiles based on studies that amplify taxonomic marker gene regions (amplicons), shotgun metagenomics studies, and shotgun metatranscriptomics studies. We selected taxonomic profiles that were constructed with the 4.1. pipeline version of MGnify and based on the small subunit (SSU) rRNA gene, contained at least 50,000 taxonomically annotated reads at the rank superkingdom, and had less than 10% of those reads classified as eukaryotic. We randomly picked one taxonomic profile per sample in cases where there were multiple. To balance the large overrepresentation of several environments in the database (for example human gut, soil, and ocean), at most 1,000 samples were randomly selected per annotated biome. The 22,518 selected samples **(Supp. table 1)** spanned 140 different annotated biomes across a wide geographical range, and consisted of amplicon, metagenomic, and metatranscriptomic studies **(Supp. fig. 1)**.

We removed eukaryotic classifications including those classified as mitochondria and chloroplast from the taxonomic profiles, and those not classified at the taxonomic rank superkingdom. When relative abundances were used, they were calculated as the number of reads assigned to a taxon divided by the total number of prokaryotic reads, unless otherwise stated in the **section ‘Ecological dissimilarity measures’**.

### Ecological dissimilarity measures

We calculated ecological dissimilarity between all sample pairs based on their taxonomic profiles (‘compositional dissimilarity’) at different taxonomic ranks using ten commonly used ecological measures: Aitchison distance, Bray-Curtis dissimilarity, Sørensen-Dice coefficient, Jaccard distance, weighted Jaccard distance, Kendalls τ_*b*_ coefficient, Pearson correlation coefficient, Spearman’s rank correlation coefficient, unweighted Unifrac distance, and weighted Unifrac distance. Some are true distance or dissimilarity measures, whereas others can be readily converted to a scale from 0 to 1, with 0 being compositionally similar. The three correlation measures were converted to dissimilarity with the formula 0.5 − (coefficient/2), and we used 1 − Sørensen − Dice coefficient.

Taxa that were represented by less than 5 reads were removed before dissimilarity calculations. This was done per rank, and therefore the total number of included reads for a sample could differ depending on the rank considered. To ensure that the pairwise calculations are based on the deepest attainable resolution, we decided for a low absolute read cut-off as opposed to a relative abundance cut-off. For each pairwise calculation, we only included taxa that were present in the union of the two samples, thus avoiding the vast scarcity (i.e., presence of zeros in the abundance matrix) often associated with microbiome studies. This scarcity is especially likely because our study compares many different habitats. Those taxa that were only present in one of the samples were given an abundance of zero in the other for all ecological dissimilarity measures except the Aitchison distance, which cannot handle zeros. For the Aitchison distance, a pseudocount was added. This pseudocount differed per pair of samples and was based on the lowest relative abundance that could be reached by an undetected taxon, namely 1 read in the sample with the highest number of taxonomically annotated reads. We defined *N*_1_ as the sum of reads represented by the taxa in sample 1, and *N*_2_ as the sum of reads represented by the taxa in sample 2, with *N*_1_ ≥ *N*_2_. A pseudocount of 1 read was added to all taxa in sample 1, and of 1/*N*_1_ × *N*_2_ reads in sample 2.

Since the MGnify taxonomic annotation pipeline depends on sequence similarity to a reference database and environmental sequencing studies contain both known and unknown taxa ^74^, many reads in a sample are not classified at all ranks (superkingdom, phylum, class, order, family, genus, species). Furthermore, taxonomy is incomplete, with lower rank classifications sometimes present but intermediate ones missing. For example, taxonomy of the phylum *Cyanobacteria* is debated ^75^ and currently only one order has a taxonomic annotation at rank class in NCBI taxonomy. Likewise, the genus *Methyloceanibacter* of the order *Rhizobiales* does not have a family annotation yet ^76^. For these reasons, we calculated the ecological dissimilarity measures at all ranks up to phylum with 3 different methods for dealing with unknowns in the data. For the UniFrac distances we used a different method **(see below)**. For the first approach (i), we considered any taxon on the specific rank. If there was no classification at that rank but the taxon contained lower rank classifications, the first classified rank below was used. If there was no classification at the specific rank and no lower rank classification, we used the first classified rank above. For the second approach (ii), we exclusively considered taxa that were classified at the specific rank. Taxa that were classified at lower or higher ranks alone were removed. For the third approach (iii), we treated taxa that were not classified at the specific rank but did have lower rank classifications the same as in (i). If taxa had no classification at the specific rank or at a lower rank, we used the first classified rank above, unless the taxon was present in both samples. In this case the taxon was removed. The rationale is that for these taxa it is unknown if they are the same or different for the rank of interest.

UniFrac distance takes relatedness between taxa into account. We used distance across the taxonomic tree as measure for relatedness, with distance between successive ranks defined as 1. We used the EMDUnifrac implementation ^77^, which is suited for samples with many unknowns because it allows for the placement of taxa at different ranks in the tree. UniFrac distances were calculated at ranks species, family, and class. For taxa that had no classification at the specific rank but did have a lower rank classification, we used an artificial classification based on the first classified rank below, ensuring uniqueness of the taxon and appropriate distance to the root. Taxa that did not have a classification at the specific rank or a lower rank were placed at the first classified rank above in the tree.

For some of the ecological dissimilarity calculations, number of reads per taxon were converted to relative abundance values by dividing by the sum of reads represented by the taxa in the sample, for example for Bray-Curtis dissimilarity calculations and the addition of pseudocounts before Aitchison distance calculations. As explained above, the taxa considered in a sample may differ per method of dealing with unknowns, and so may the relative abundance of a taxon.

Because the taxonomic profiles contain many unknowns at lower ranks, pairwise comparisons can be based on few taxa. For each rank and method of dealing with unknowns, samples were removed that did not contain any taxon at that rank. If the pairwise comparison was based on one taxon we set dissimilarity to 0. We removed samples from the correlation measures whose correlation coefficient with itself could not be calculated.

### Permutatational multivariate analysis of variance (PERMANOVA)

PERMANOVA pseudo F-statistics were calculated for all ecological dissimilarity measures with the scikit-bio v0.5.5 implementation **(http://scikit-bio.org/)**. As predefined groups we used either the annotated biomes of the samples, or their experiment types. P-values were based on 99 random permutations, and we calculated the coefficient of determination (*R*^2^) with the formula:

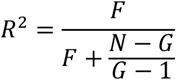

were *F* is the pseudo F-statistic, *N* the sample size, and *G* the number of groups.

### Diversity measures

Diversity measures were calculated for the subset of samples belonging to an annotated biome, and in which a taxon was found. If a subset contained less than 3 samples it was excluded from analysis. Taxa were removed whose relative abundance was less than 1 / 10,000. We used approach (ii) for dealing with unknowns as explained in the **section ‘Ecological dissimilarity measures’**.

Zeroth order alpha diversity (richness, mean number of taxa found in a set of samples) was calculated for all ranks. Zeroth, first, and second order alpha diversity (^*q*^*D*_*α*_) and beta diversity (^*q*^*D*_*β*_) were calculated on the taxonomic rank order and based on relative abundances, with ^*q*^*DD*_*β*_ defined as the total effective number of taxa (^*q*^*D*_*γ*_) divided by ^*q*^*D*_*α*_. ^*q*^*D*_*γ*_ was calculated based on summed relative abundance of the individual samples. For the first and second order diversity measures two samples were excluded that did not contain any classification at order rank after the relative abundance threshold. When the terms alpha and beta diversity are used, we refer to first order diversity measures (on the rank order) unless otherwise stated. For a more in-depth discussion of these diversity measures see ^78^. Shannon entropy and Gini-Simpson index that were used for diversity calculations were calculated with the scikit-bio v0.5.5 implementation.

### Local dominance and Shannon entropy across samples

For each taxon, we calculated local dominance and Shannon entropy. Local dominance was defined as mean relative abundance across all samples in which the taxon was found. Shannon entropy (base *e*) was used as a measure for randomness of its relative abundances across these samples (*N*), and was normalized by dividing by In (*N*).

### Social niche breadth definition

Social niche breadth (SNB) was defined as the mean of the pairwise dissimilarity between the samples in which a taxon was found, *n*:

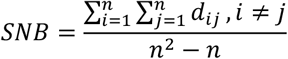

with dissimilarity *d_ij_* based on Spearman’s rank correlation coefficient (0.5 − (*ρ*/2)) on the rank order with method (ii) for dealing with unknowns, see the **section ‘Ecological dissimilarity measures’**. A taxon was considered present in a sample if it had a relative abundance of at least 1 × 10^−4^. A taxon with a low score is thus found in samples with similar taxonomic profiles (social specialist), and a taxon with a high score in dissimilar samples (social generalist). Taxa that were present in less than 5 samples were removed from analyses, unless otherwise stated.

To benchmark SNB, we also calculated SNB with different detection thresholds of 1 × 10^−3^ and 1 × 10^−5^. Social niche breadth was moreover calculated for imaginary taxa (iSNB) that were present in all samples from a given annotated biome, in half of the samples from a given annotated biome (100 random permutations per annotated biome), and in all samples from pairs of annotated biomes. In addition, iSNB was calculated for randomly picked sets of samples of equal size to the encountered taxa (100 random permutations per sample size). Lastly, we calculated SNB for real taxa only based on the marine and human hierarchical subsets of the samples.

High-ranking taxa whose genera where significantly (p < 0.05) biased towards high or low SNB were decided based on the Mann-Whitney *U* test of the distribution of its genera versus the distribution of all other genera.

### Selection of genomes

We downloaded all genomes from the Pathosystems Resource Integration Center (PATRIC) genome database ^62^ that had a quality marked as ‘good’ and were not plasmids on 14 November 2019. We only included genomes for which we had a valid taxonomy id in our NCBI taxonomy ^79^ files that were downloaded on the same date. PATRIC contains identical genomes with different identifiers. We identified replicate genomes based on concatenated DNA sequence and concatenated sorted DNA sequence, and removed all but one. In cases where identical genomes had different taxonomic taxa, all were discarded.

Completeness and contamination estimates were generated with CheckM v1.0.7 ^80^ in the lineage-specific workflow. We excluded genomes whose *completness* − 5 × *contamination* < 70. The final selection consisted of 225,101 prokaryotic genomes representing 34,304 species, from both cultures and environmental sequencing projects **(Supp. table 4)**.

Inferences about (pan) genome size and genomic functions **(see below)** and number of subtaxa were made at all taxonomic ranks, and reconstructions at higher ranks were based on lower rank taxa in the PATRIC database that were not always present in the MGnify dataset.

### Genome size estimates and GC content

Genome size in number of base pairs and number of coding sequences (CDS), and GC content were obtained from the metadata in the PATRIC database. We corrected genome size estimates by taking completeness and contamination into account, via multiplication with a scaling factor s:

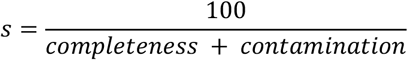

For each measure including the number of CDS per million base pairs we reconstructed species by averaging the values of its genomes. For higher ranks, mean values were calculated for all species belonging to the taxon. Some high-ranking taxa contain many low-ranking taxa from the same taxonomic group such as genus or family. To correct for this overrepresentation and possible skew towards the values of these taxa, we also calculated mean values by averaging over the taxonomy at all ranks (‘taxonomy corrected’ values). For example, the size of a family is the mean of the sizes of its genera, which are the mean of the sizes of its species. These values were calculated for the ranks family and higher and can be found in **Supp. table 4**.

### Genome functions

Functional profiles of the genomes were created based on the PATRIC annotations of coding sequences for three functional universes: subsystems, GOs, and pathways. For GOs the profiles were based on the exact terms found in the annotation files, and for subsystems we made different profiles for the ‘name’ and ‘subclass’ level of the hierarchy. Genomes with ≤ 20 unique functions were discarded from further analyses for subsystem names, GOs, and pathways. For the included list of genomes in each analysis see **Supp. table 4**.

Functional genome size was defined as the number of unique functions present in a species. A function was considered present in a species if at least 50% of its genomes contained it. Mean functional genome sizes were calculated for all taxa, as well as the standard deviation. Pan genomes were defined at all ranks as the total set of unique functions present in the genomes of a taxon.

Pan genomes can be open or closed, meaning they can be more or less susceptible to changes in gene content ^60,61^. We devised a score that represents pan genome openness for all ranks higher than species. Pan genome openness was defined as the total pan genome size divided by the mean pan genome size of a species. Because taxa with many daughter taxa tend to have large pan genomes, we also calculated pan genome features for a random subset of 3 daughter species (1,000 random permutations per taxon) to correct for this effect of taxonomy. This measure thus reflects how many functions are on average added to the pan genome by including two more species. Permuted measures were calculated for all taxa with at least 3 species.

### Gene set enrichment analysis (GSEA) of pan genomes

To detect functions that were significantly enriched in social specialists and generalists, we deployed gene set enrichment analysis (GSEA) ^71^ based on the pan genomes of genera. We performed a GSEA on all genera sorted by SNB to compare specialists to generalists, and on specialists genera (SNB < 0.35) sorted by alpha diversity to compare low diversity specialists to high diversity specialists. We used the classical Kolmogorov-Smirnov statistic for the enrichment score (*p* = 0). Enrichment score normalisations and p-values were based on 100,000 random permutations of the gene set. Multiple hypothesis correction was carried out via the false discover rate (FDR) as suggested in ^71^. GSEA computations were done with a modified version of the algorithm.py script from GSEApy v0.7.3.

### Growth rate and clade age estimates

We downloaded the maximal growth rate predictions of RefSeq genomes from the EGGO database ^54^, and defined species and genus rank maximal growth rates as the mean of its genomes.

Clade ages were based on the TimeTree database ^66^. Times to the first and last common ancestor were extracted from the species rank phylogenies of *Bacteria* and *Archaea* using ete3 v3.1.1 ^81^.

### Software packages used for calculations and visualizations

Calculations were done with the Python 3 standard library, NumPy ^82^, and the SciPy library ^83^, unless otherwise stated. Visualisations were done with Python 3 and Matplotlib ^84^ in JupyterLab **(https://jupyter.org/)**, with the use of NumPy, pandas ^85^, and seaborn **(https://seaborn.pydata.org/)**. Principal coordinates analysis (PCoA) was performed with the scikit-bio v0.5.5 implementation. t-distributed stochastic neighbour embedding (t-SNE) was performed with scikit-learn v0.21.3 ^86^. Samples were drawn on the world map with Cartopy v0.17.0 **(http://scitools.org.uk/cartopy)**. The taxonomic tree was visualised with iTOL ^87^, and the hierarchical tree of annotated biomes with ete3 v3.1.1 ^81^.

## Supporting information

Supplementary results and discussion

