## Supplementary results and discussion for "On specialists and generalists: niche range strategies across the tree of life"

#### A cross-biome dataset

We compiled a diverse set of 22,518 environmental sequencing samples from 592 studies, spanning 140 annotated biomes across a wide geographical range based on the MGnify resource [1] **(Supp. fig. 1, Fig. 1, Suppl. table 1, see Methods for selection criteria)**. MGnify uses standardised pipelines to process environmental sequencing datasets, allowing for the comparison of samples across a wide range of different environments, studies, and datatypes. We only included datasets that were processed with the MGnify version 4.1 pipeline.

As expected, samples from similar annotated biomes clustered together based on microbial composition, despite the samples coming from vastly different locations and study designs **(Fig. 1d, Supp. fig. 2a, Supp. fig. 3)**. Samples were mostly separated by association to a vertebrate host versus free-living habitats, and saline versus non-saline habitats [2–5] **(Supp. fig. 4a, Supp. fig. 2b)**. An exception are fish, whose foregut and intestinal microbiomes were more similar to microbiomes from aquatic habitats than to those in other vertebrate guts **(Supp. fig. 3)**. Within free-living habitats, saline samples differed from non-saline samples including soils. Aquatic sediments resembled their saline or non-saline provenance. In line with earlier findings [3], invertebrate-associated samples clustered together with free-living samples and not with vertebrate-associated samples. Invertebrates in our dataset include sponges, molluscs, *Cnidaria*, and *Echinodermata* whose internal microbiomes are in direct or semi-direct contact with the surrounding environment, and marine arthropods like *Calanus finmarchicus* **(Supp. table 1)**.

Host-associated samples typically had lower taxa richness and alpha diversity than free-living samples **(Supp. fig. 4b,c, Supp. fig. 2c-e, Supp. fig. 5a)**. Rhizospheres have been shown to resemble soils in terms of richness [4] and we also observed this **(Fig. 1e)**. Notably, annotated biomes with a high alpha diversity had a low beta diversity, while annotated biomes with low alpha diversity had either low or high beta diversity **(Fig. 1f)**.

While taxa richness increases towards low taxonomic ranks, richness in the microbiomes was lower at the species rank and in some cases also at the genus rank than at higher ranks **(Fig. 1e)**. This anomaly reflects the still low classification rate of the organisms in natural environments at the species and genus ranks. In addition, low rank classifications more easily fall below our detection limit of 1 / 10,000 than a higher rank classification whose abundance is the sum of its lower ranks.

#### Quantifying microbial niche breadth

To quantify the range of habitats in which a microbial taxon is found we formulated a niche breadth score that is data-driven and independent of human-defined biome annotations. To do this, we calculated the dissimilarity between taxonomic profiles of pairs of samples (see below), and defined the "social niche breadth" (SNB) of a taxon as the mean pairwise dissimilarity between the microbial communities of the samples where it is found. Thus, taxa that always occur in samples with similar microbial composition have a low SNB ("social specialists"), whereas taxa that occur in dissimilar samples have a high SNB ("social generalists").

#### Benchmarking microbiome dissimilarity measures

To arrive at a quantitative niche breadth definition that optimally reflects annotated biomes as recognised by the research community, we benchmarked 150 different microbiome dissimilarity measures for their correspondence with the biome annotations of the underlying datasets. Many dissimilarity measures have been proposed in ecological literature, each with their own merit. For example, the Aitchison distance [6] is relevant for microbiomes because it takes the inherent compositionally of sequencing data into account [7], while the Unifrac distance considers phylogenetic (or taxonomic) information [8]. Measures that take relative abundances into account (weighted measures) better reflect quantitative relationships between taxa than unweighted measures, but put less emphasis on low abundant taxa that might be instrumental for ecosystem functioning [9]. Other factors of importance include the taxonomic rank of comparison, as well as the method for handling unknowns (sequences that cannot be classified).

We calculated ten ecological dissimilarity measures between all 253,518,903 sample pairs at six taxonomic ranks and with four different methods for dealing with unknowns **(see Methods)**, totalling 150 different measures **(Supp. fig. 6a-d)**. Based on a comparison to annotated biomes using PERMANOVA, we found that these groups are best represented by a dissimilarity measure based on an inverted Spearman's rank correlation coefficient $(0.5 - (\rho/2))$ at the taxonomic order rank while ignoring unknowns **(Supp. fig. 6a-d)**. Notably, performing the same analysis with experiment type (amplicon, metagenomic, metatranscriptomic, or unknown) as the predefined groups showed very low $R^2$ values **(Supp. fig. 6a-d)**, implying a low impact of the study design on the ecological clustering and justifying our decision to include these different data types in this global analysis. We thus use the Spearman-based microbiome dissimilarity score to quantify SNB, while noting that another choice would not qualitatively affect our results, as niche breadth scores based on six alternative ecological dissimilarity measures showed a high correlation with the one we selected **(Supp. fig. 6e)**. In addition, we investigated the robustness of our results to our choice for the mean pairwise dissimilarity by comparing it to the median and third quartile, and found high correlations ($\rho = 0.977$ (p: 0.000) and $\rho = 0.967$ (p: 0.000), respectively; **Supp. fig. 7)**. In agreement with our premise that social niche breadth should reflect the cooccurrence of a taxon with other taxa, our SNB score is strongly negatively correlated with the fraction of shared taxa between samples ($\rho$ = -0.878 (p: 0.000); **Supp. fig. 8)**.

#### Social niche breadth robustly reflects community heterogeneity

To investigate the robustness of the SNB score to sampling bias, we further calculated social niche breadth for imaginary taxa (iSNB) that occur in all samples of an annotated biome. This showed a strong association between iSNB and the beta diversity of an annotated biome **(Supp. fig. 9a, Supp. fig. 4d)**. Thus, taxa that are ubiquitously present in a very heterogeneous annotated biome would have a high SNB, while taxa that are ubiquitous in an annotated biome with a low turnover rate would have a low SNB. Importantly, iSNB does not depend on the number of available samples for a given annotated biome **(Supp. fig. 9b)**. Random subsets of samples from single annotated biomes showed low variation in iSNB **(Supp. fig. 9c, Supp. fig. 4e)**, implying robustness of SNB to sporadically missed presence of a taxon in a sample, but standard deviation increased when the number of samples becomes very low **(Supp. fig. 9c, Supp. fig. 4e)**. For this reason, we use caution when interpreting SNB of rare taxa, i.e. that are present in only a few samples, and exclude taxa that are present in less than 5 samples from our analyses. iSNB calculated for imaginary taxa that occur across two annotated biomes revealed low iSNB for highly similar annotated biomes like different human oral sites **(Supp. fig. 10**, top left corner**)**. In addition, even though presence in a single annotated biome with low beta diversity results in a low iSNB, presence in two different annotated biomes with low beta diversity still results in a high iSNB if they are very different from each other **(Supp. fig. 10)**.

For real microbial taxa, we observed a striking independence of SNB on the number of samples in which a taxon is found **(Fig. 2a-c)**. Some moderately ubiquitous taxa are exclusively present in similar samples (low SNB), whereas many uncommon taxa are present in very different samples (high SNB). That some specialist taxa are still quite ubiquitous can partly be explained by the overrepresentation of some environments in our microbiome dataset, even though we selected a maximum of 1,000 samples per annotated biome **(Supp. fig. 1b)**. Further, we observed widely different SNB for taxa encountered in the same number of annotated biomes **(Fig. 2b)**, pointing to differences in turnover rate between annotated biomes and heterogeneity within annotated biomes as discussed above. Nonetheless, many taxa that are found in only a few samples have a low SNB **(Supp. fig. 11a)**, indicating that the samples where they are found are similar in composition, and suggesting that rare taxa are often specialists. The most cosmopolitan taxa are all generalists (high SNB) **(Fig. 2a-c)**, since they are present in many dissimilar samples.

There is a good correlation between SNB of a taxon and its SNB calculated based only on the human and marine subsets of samples ($\rho = 0.546$ (p: 0.000) and $\rho = 0.662$ (p: 0.000), respectively; **Supp. fig. 12)**, implying that the sampling of annotated biomes does not strongly affect our calculated SNB values and suggesting that our general results would be qualitatively similar if different habitats were sampled.

The detection limit of taxa in environmental sequencing datasets is an important parameter that could influence SNB, as higher detection thresholds obscure our view of rare taxa [9] and decrease the number of samples and habitats in which taxa are found. To assess this effect, we calculated SNB with a ten-fold higher ($1 \times 10^{-3}$) and ten-fold lower ($1 \times 10^{-5}$) detection threshold than used for our main results **(Supp. fig. 13)**, and observed shifts in SNB as expected; overall, taxa become more specialist with a higher and more generalist with a lower detection threshold. Importantly, the list of taxa ranked by SNB was consistent **(Supp. fig. 13c,g)**, especially if uncommon taxa were excluded **(Supp. fig. 13d,h)**. In addition, exclusion of taxa that have very low relative abundance across samples does not change the distribution of SNB **(Supp. fig. 11b)**.

Together, the results presented in this section show that the SNB score is robust to the specific community dissimilarity measure, sampling bias, and detection threshold, suggesting that our results represent a meaningful quantification of microbial niche range.

### Supplementary figures

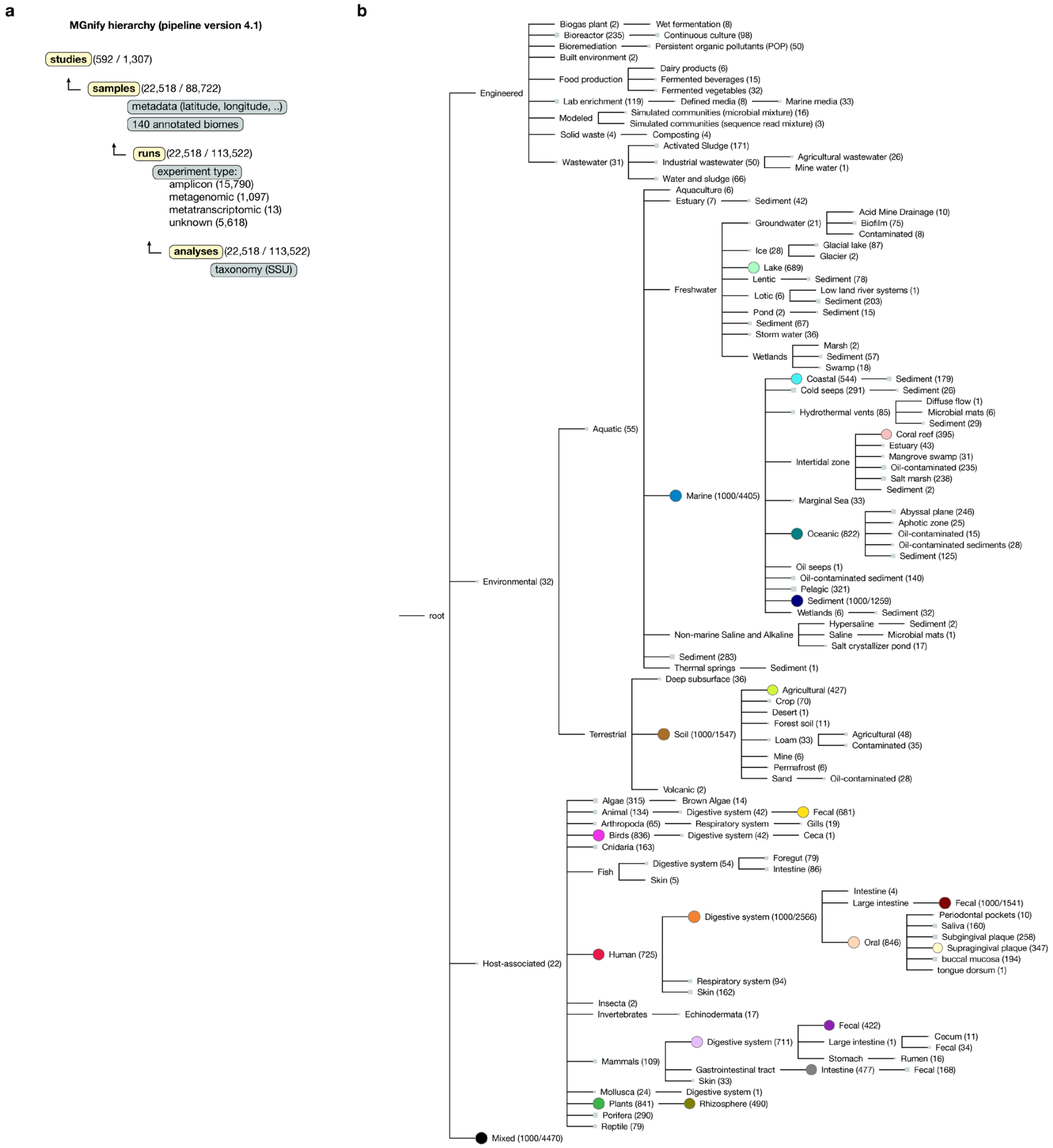


**Supplementary figure 1. Selection of samples and annotated biomes from the MGnify resource.** (**a**) Studies in MGnify are divided into samples, which have associated runs and taxonomic analyses. Numbers within brackets indicate the number of selected instances out of the total in the resource that is annotated with pipeline version 4.1. A single taxonomic analysis on SSU level was picked per sample. For other selection criteria see **Methods**. (**b**) Hierarchical tree of annotated biomes. Numbers within brackets and size of markers indicates the number of samples from the annotated biome. If more than 1,000 samples met the selection criteria, 1,000 samples were picked at random and the total sample pool is indicated as the second number. Colours correspond to colour-coding of biomes in **Fig. 1**.

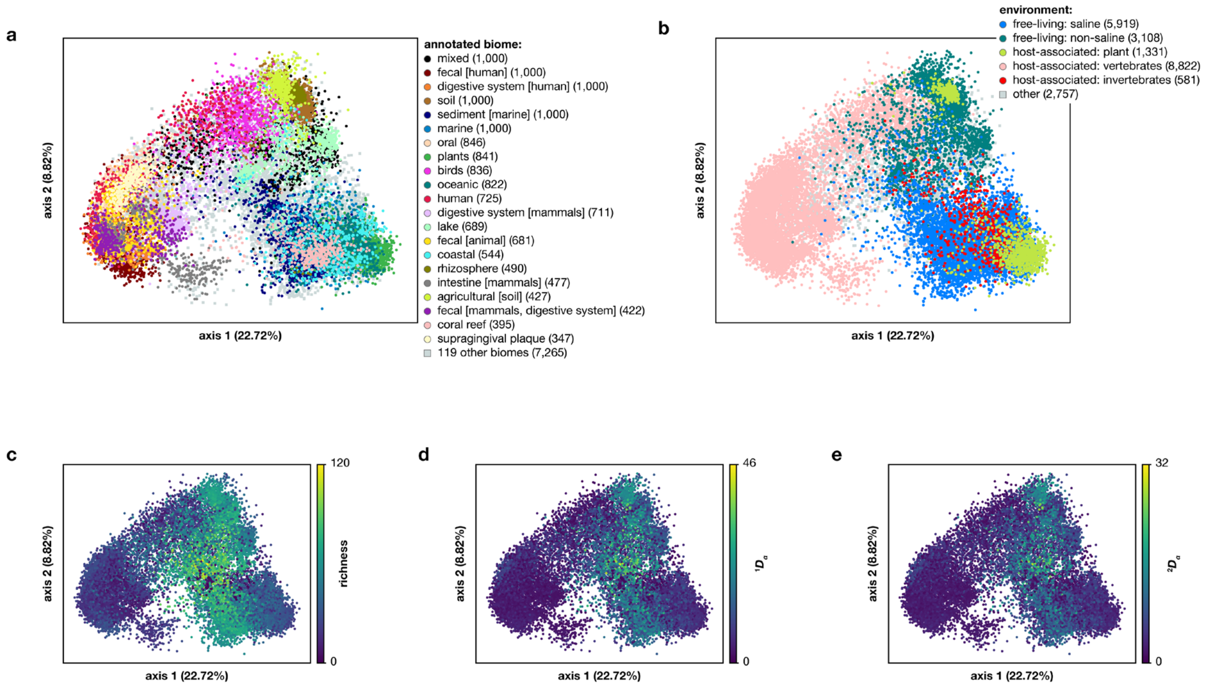


**Supplementary figure 2. PCoA visualisation of 22,518 microbiomes.** (**a**) Samples from similar annotated biomes cluster together based on taxonomic profile, with the same ecological dissimilarity measure used as for SNB, namely Spearman's rank correlation coefficient $(0.5 - (\rho/2))$ of known taxa at rank order. (**b**) Samples are separated by host-association and salinity. Invertebrate-associated communities cluster together with free-living communities and not with vertebrate-associated communities. See **Supp. fig. 3** for the division of annotated biomes in free-living and host-associated. See **Supp. fig. 4a** for a t-SNE visualisation of the same data. (**c-e**) Alpha diversity of samples on the rank order for three different diversity measures, (c) zeroth order diversity (richness), (d) first order diversity ($e^{Shannon\ index}$), and (e) second order diversity (inverse Simpson index).

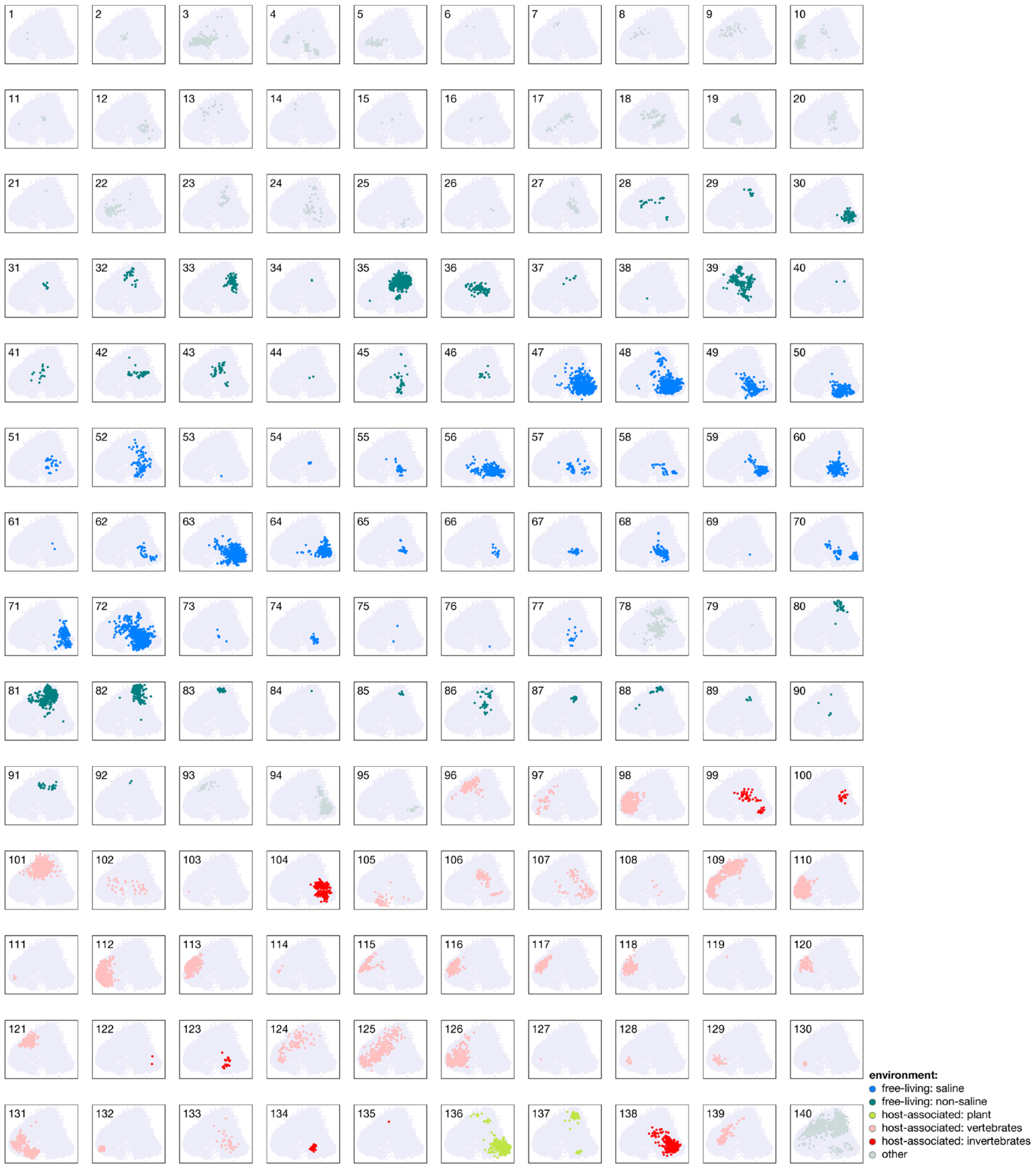


**Supplementary figure 3. Location of all annotated biomes on the PCoA.** Annotated biomes are coloured according to their environment type, free-living or host-associated. The 'Animal' biomes contain samples from frog, iguana, cats, dog, pigeon, fish, rat, cow, pig, mouse, poultry, and mammalia-associated habitat **(Supp. table 1)**, and are thus included in the vertebrate environment type.

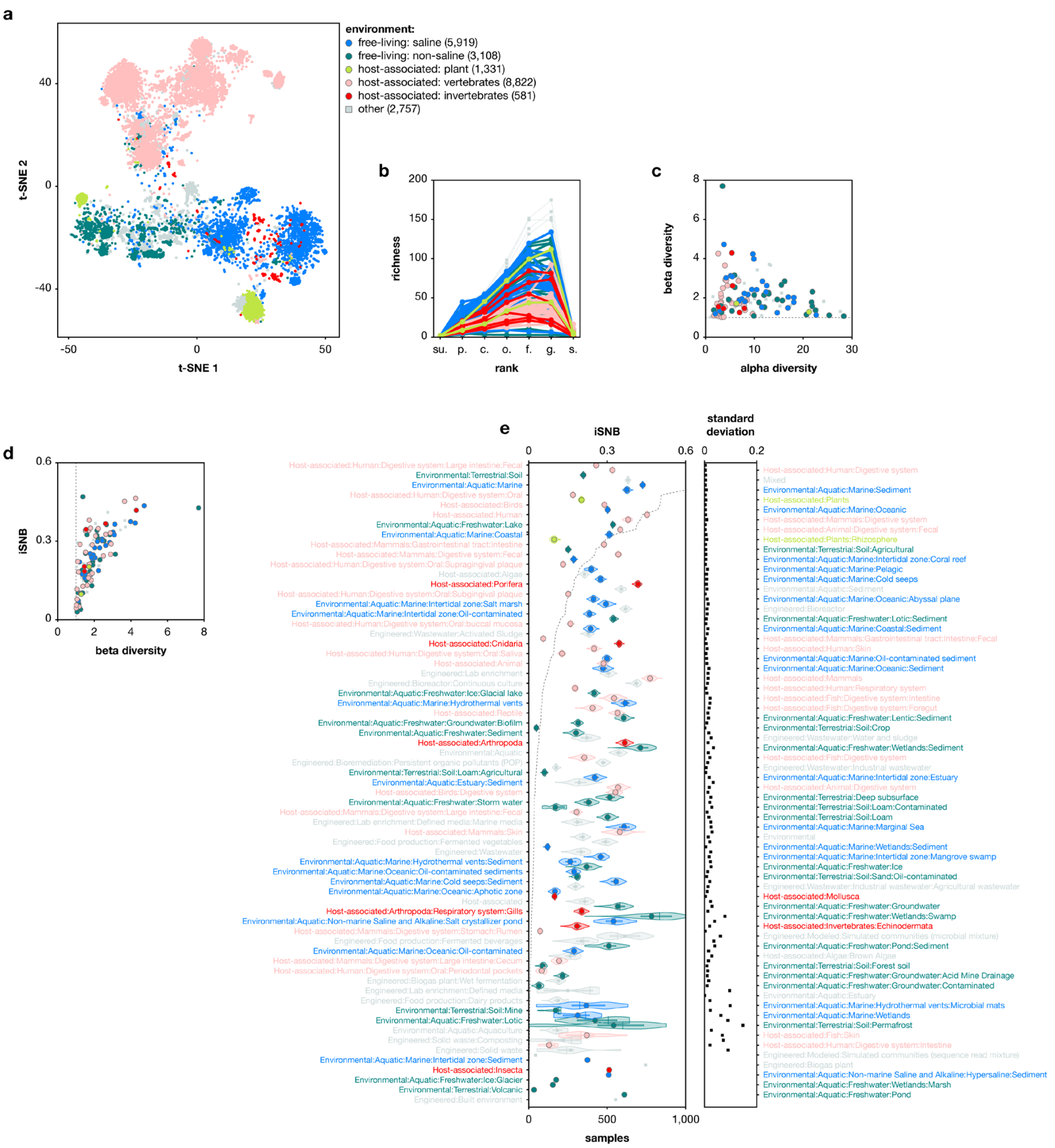


**Supplementary figure 4. Remake of some figures colour-coded according to environment type.** (**a**) Idem to Fig. 1d. (**b**) Idem to Fig. 1e. (**c**) Idem to Fig. 1f. (**d**) Idem to Supp. fig. 9a. (**e**) Idem to Supp. fig. 9c.

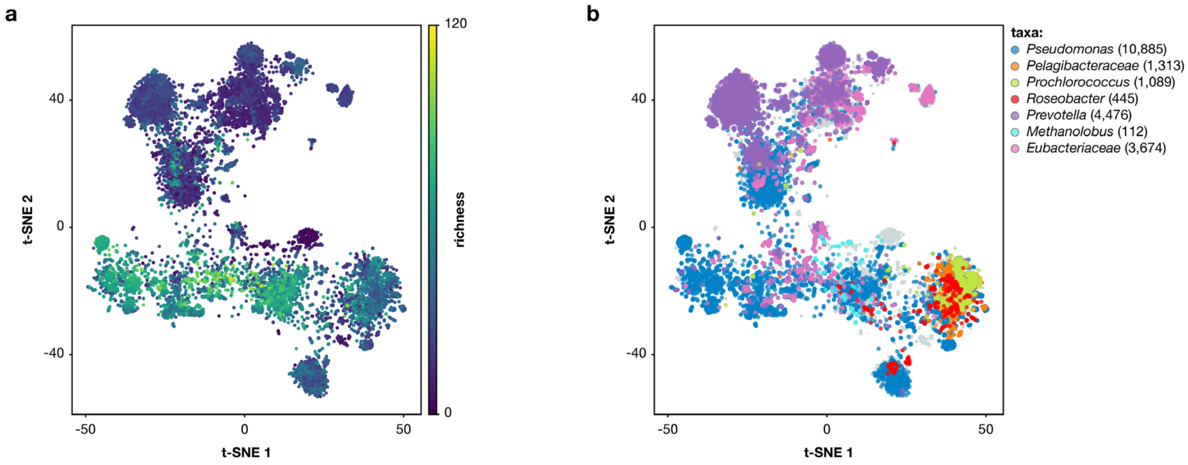


**Supplementary figure 5. Richness of samples and presence of some taxa visualised on the t-SNE of Figure 1d.** (**a**) Zeroth order alpha diversity (richness) on the rank order of samples. (**b**) Presence of some microbial taxa.

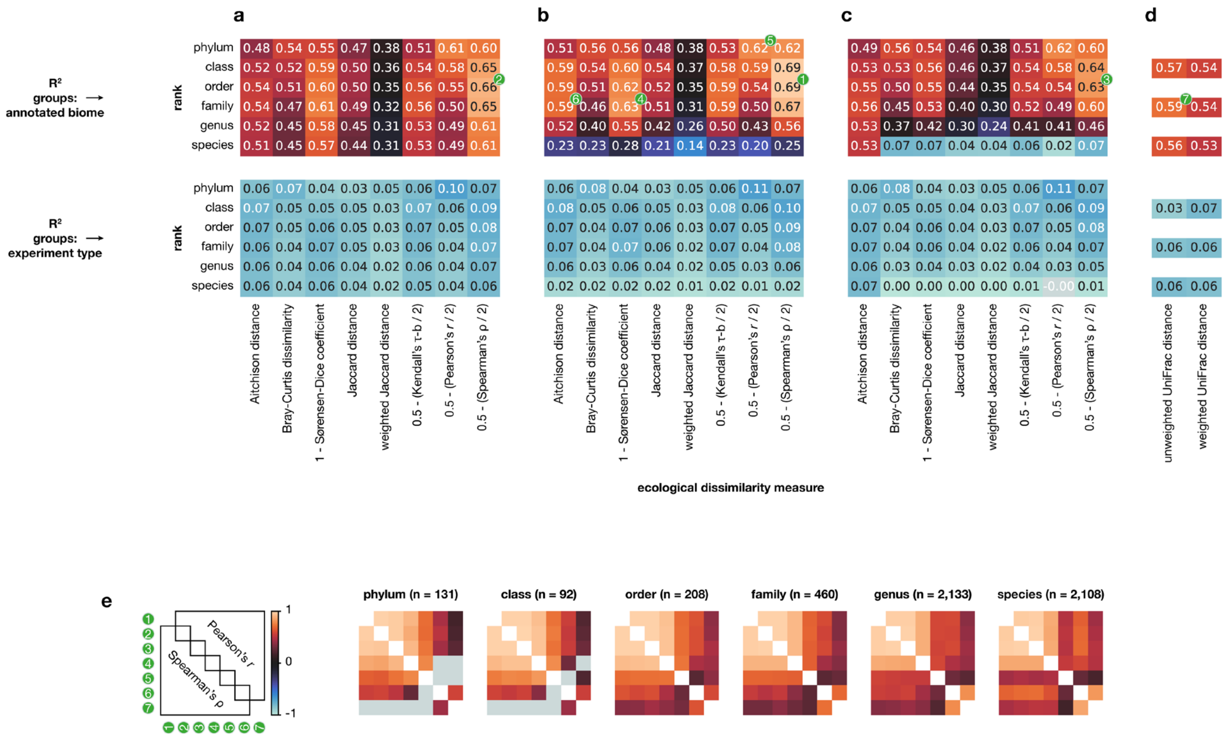


**Supplementary figure 6. Selection of ecological dissimilarity measure.** PERMANOVA results with annotated biomes or experiment type (amplicon, metagenomic, metatranscriptomic, or unknown) as predefined groups (see **Supp. fig. 1**). Dissimilarity between any two samples was calculated for 10 different dissimilarity measures and 3 different methods for dealing with unknowns. See **Methods** for a description of these methods. (**a**) Method i. (**b**) Method ii. (**c**) Method iii. (**d**) Unifrac distances. (**e**) Correlations between niche breadth of taxa calculated with 7 different ecological dissimilarity measures (green badges in panels a-d) at different taxonomic ranks. Values in panels a-e that are not significant ($p > 0.05$) are coloured grey.

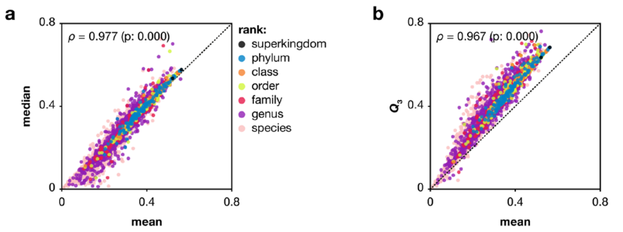


**Supplementary figure 7. SNB is robust against the choice for mean pairwise distance between the samples containing a taxon.** (**a**) SNB of taxa calculated with mean versus median pairwise distance. (**b**) SNB of taxa calculated with mean versus the 75th percentile pairwise distance. Spearman's rank correlation coefficient is indicated.

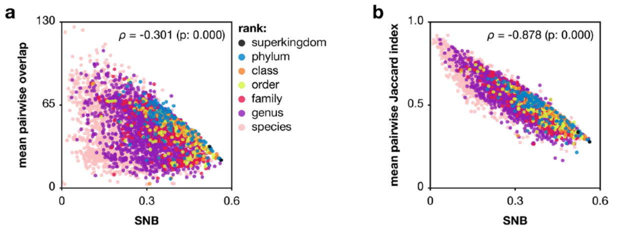


**Supplementary figure 8. SNB correlates with the fraction of overlapping taxa between the samples containing a taxon.** (**a**) SNB of taxa versus mean absolute pairwise overlap. (**b**) SNB of taxa versus mean relative pairwise overlap. Overlap was calculated on the taxonomic rank order. Spearman's rank correlation coefficient is indicated.

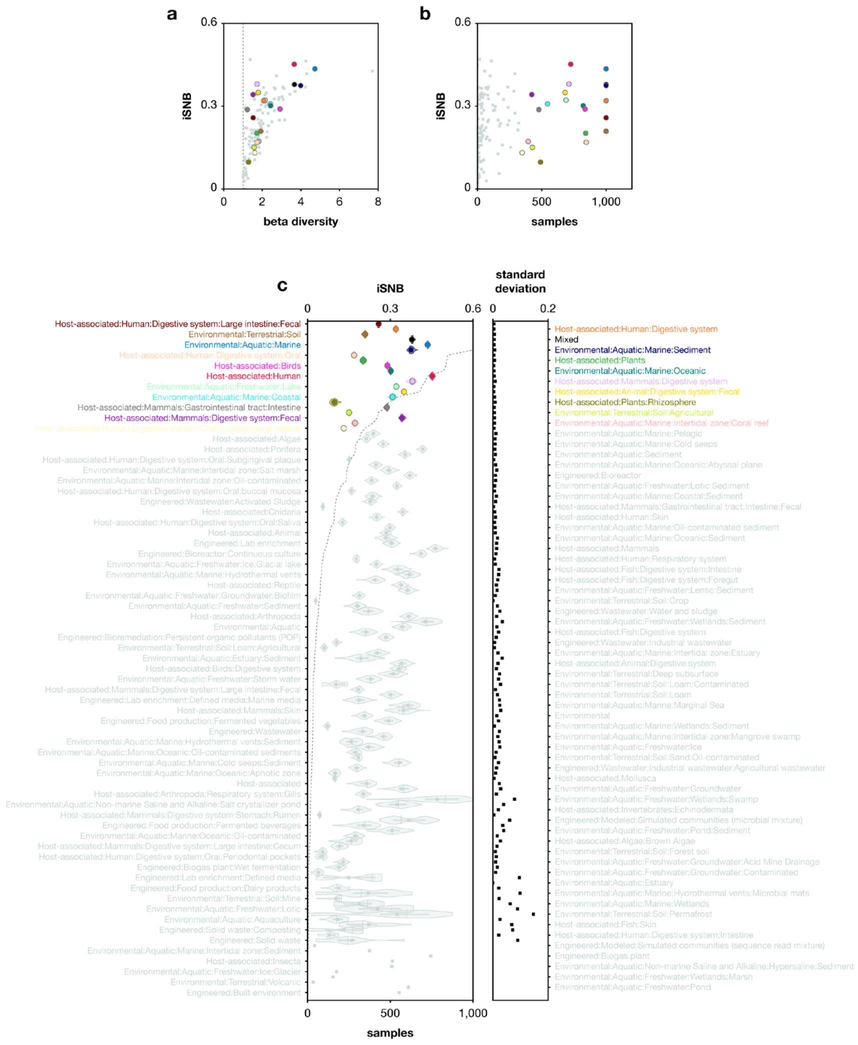


**Supplementary figure 9. SNB reflects beta diversity and is largely independent of the number of samples in which a taxon is detected.** (**a**) Beta diversity of annotated biomes versus niche breadth for imaginary taxa (iSNB) that are found in all samples of an annotated biome. (**b**) Number of samples of annotated biomes versus their iSNB. (**c**) iSNB calculated for hypothetical taxa that are found in all samples of an annotated biome (dots) and 100 times randomly picked 50% of the samples (violins). Annotated biomes are sorted according to number of samples (dashed line). The standard deviation of iSNB of the 100 randomly picked samples is depicted on the right.

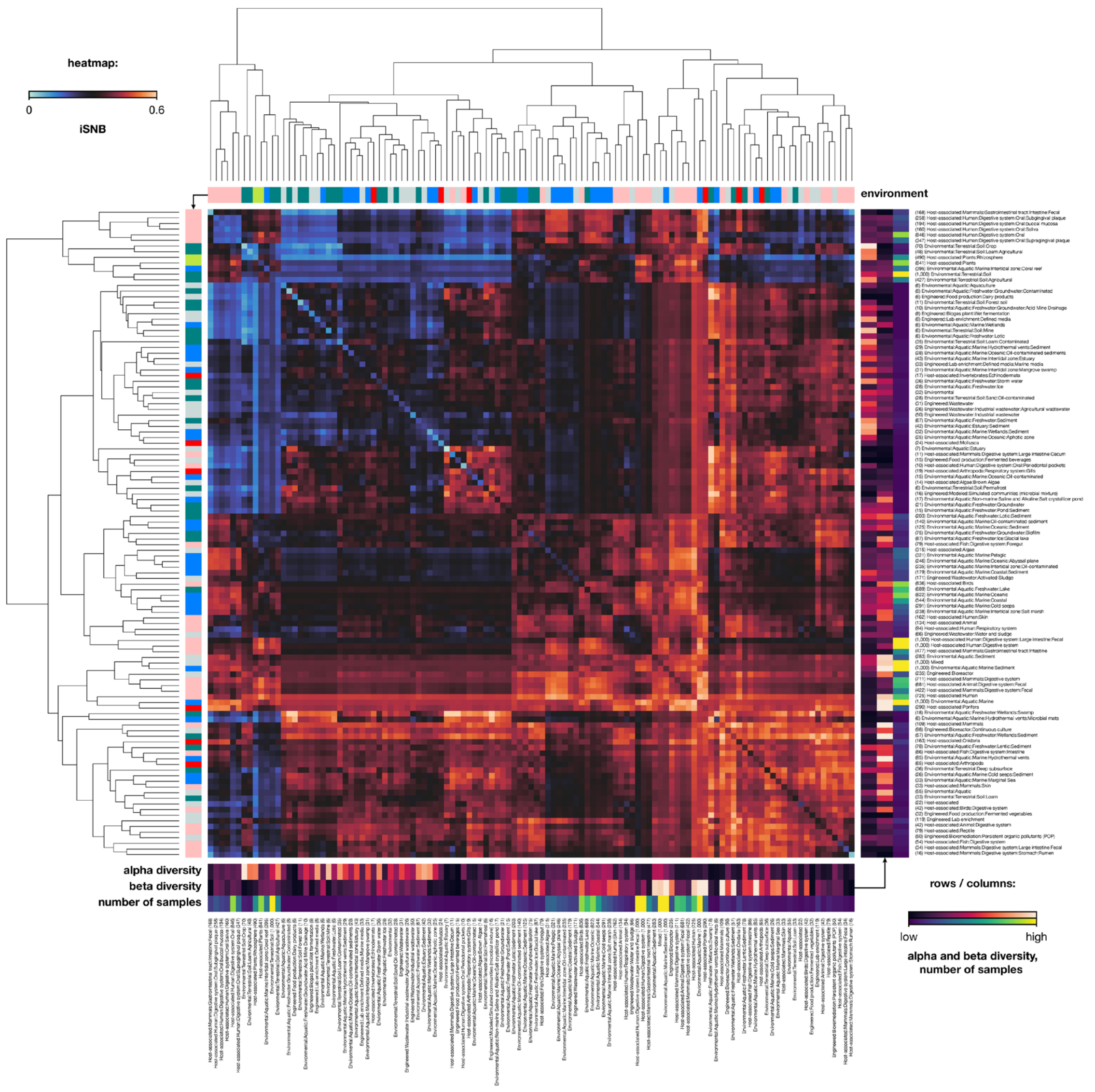


**Supplementary figure 10. Niche breadth of hypothetical taxa (iSNB) that are present in all samples of all combinations of two annotated biomes.** Hierarchical clustering of the heatmap is based on euclidean distance and the UPGMA algorithm. Alpha and beta diversity and number of samples are indicated with colour-coding along the axes. Number of samples are also indicated in the labels within brackets. The environment type colour-coding corresponds to **Supp. fig. 3**.

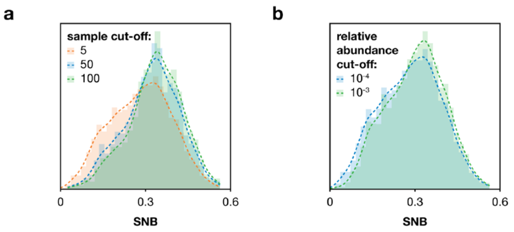


**Supplementary figure 11. Response of the overall distribution of SNB scores to different parameter cut-offs in our pipeline.** (**a**) The minimum number of samples in which a taxon must be found. (**b**) The minimum relative abundance that must be reached by a taxon in at least 1 sample.

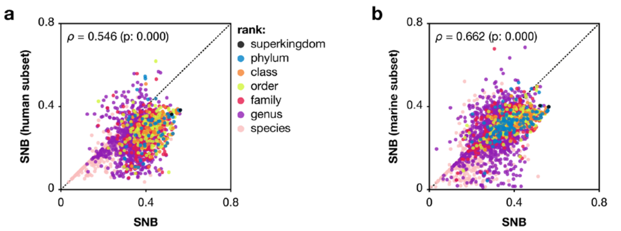


**Supplementary figure 12. SNB is relatively invariant to environmental scale.** SNB based on all samples versus SNB based on the hierarchical subsets of (**a**) human annotated biomes and (**b**) marine annotated biomes. Spearman's rank correlation coefficient is indicated. See **Supp. fig. 1b** for which annotated biomes are included in the 'Human' and 'Marine' hierarchical subsets.

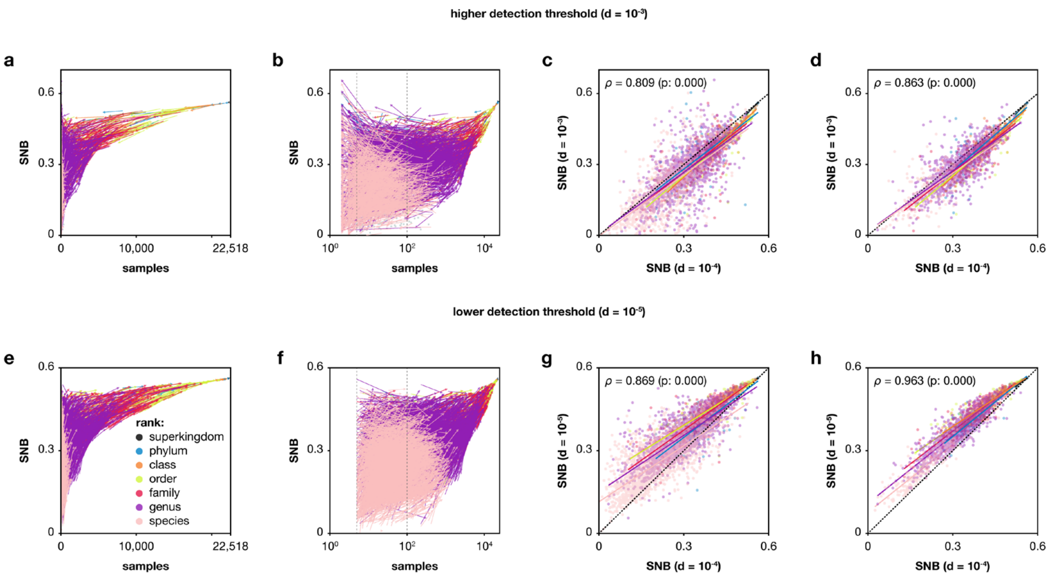


**Supplementary figure 13. Effect of the detection threshold on SNB.** (**a,e**) Arrows show the change in number of samples and SNB due to a higher and lower detection threshold. (**b,f**) The same figure but with a logarithmic x-axis. Dashed lines indicate a presence in 5 samples which is the default cut-off for taxa to be included in most analyses in this study, and a presence in 100 samples which is the cut-off for panels d and h. (**c,g**) SNB with default detection threshold versus SNB with a higher or lower detection threshold. Coloured lines are linear regression lines for different taxonomic ranks. Taxa that had a presence in less than 5 samples with a higher detection threshold (see panel b) are included. (**d,h**) The same figure but with taxa that are present in less than 100 samples removed (dashed lines in panels b and f). Spearman's rank correlation coefficient is indicated.

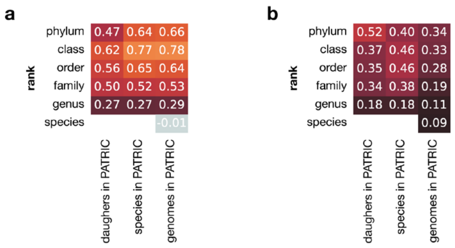


**Supplementary figure 14. Correlation between SNB of a taxon and its number of subtaxa in the PATRIC database.** Daughters are the number of taxa one rank below the current rank. (**a**) Spearman's rank correlation coefficient, and (**b**) Pearson correlation coefficient. The correlation with number of genomes in PATRIC on the species rank in panel a was not significant ($p > 0.05$) and is coloured grey.

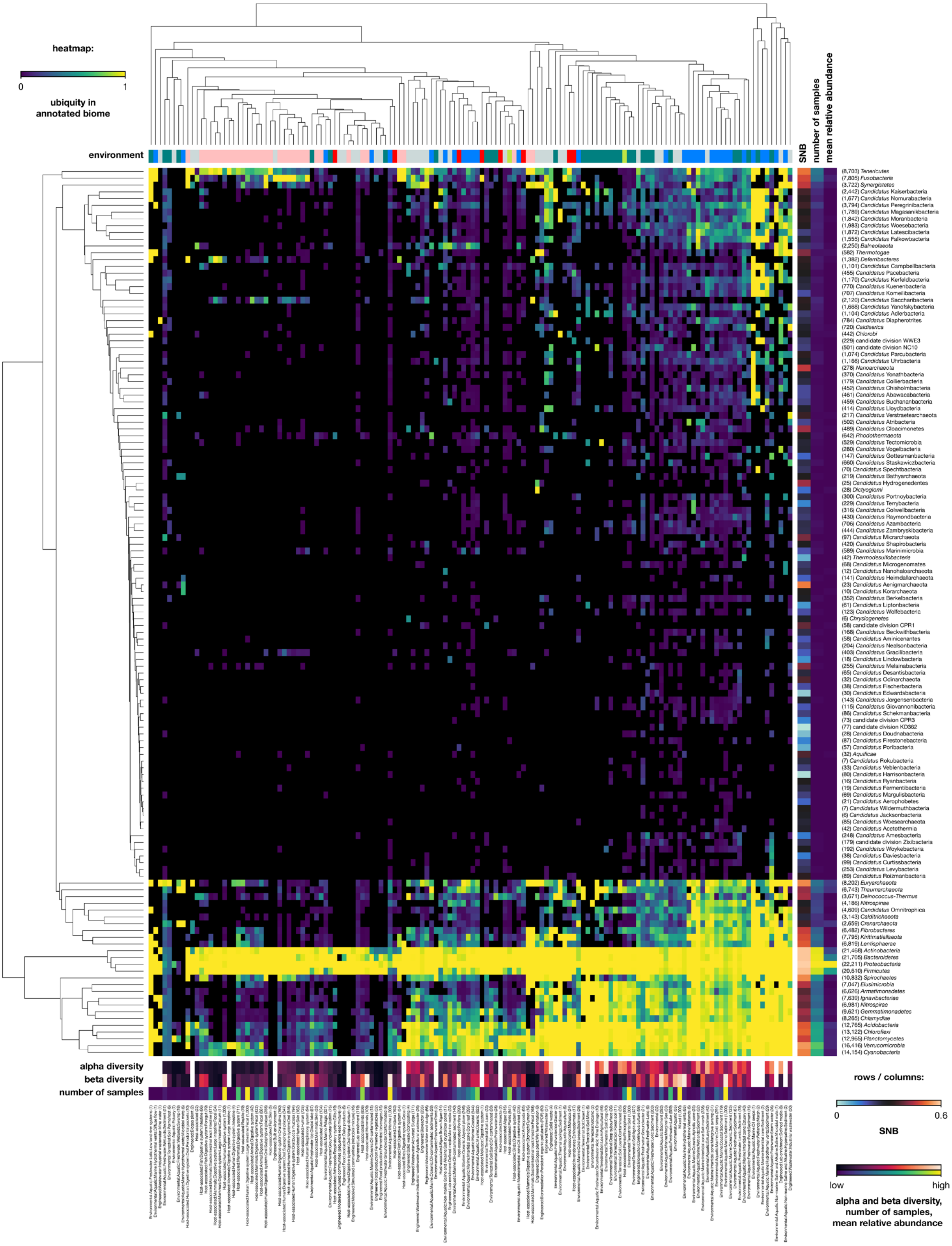


**Supplementary figure 15. Distribution of phyla in annotated biomes.** Heatmap shows the fraction of samples of an annotated biome in which the phylum is found. Hierarchical clustering of heatmap is based on Euclidean distance and the UPGMA algorithm. Alpha and beta diversity, and number of samples of biomes are indicated with colour-coding along the x-axis. SNB, number of samples, and mean relative abundance of a phylum are indicated with colour-coding along the y-axes. Number of samples are also indicated in the labels within brackets. The environment type colour-coding corresponds to **Supp. fig. 3**.

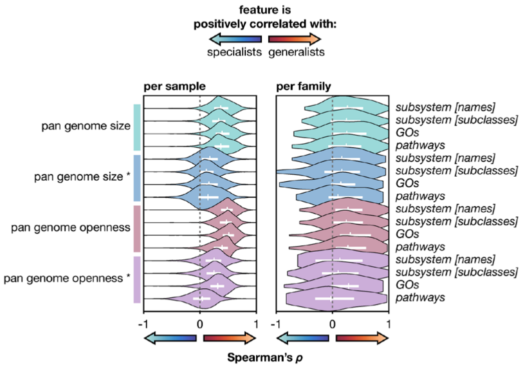


**Supplementary figure 16. Correlations between SNB and pan genome size and pan genome openness do not depend on a higher number of species in generalists.** $\rho$ per sample and per family between SNB and pan genomic features on the rank genus. Violins depict the distribution of $\rho$ across all communities or across all families with at least 5 genera. Measures are in number of unique functions for the functional universe on the right. Measures with an asterisk (*) are based on the mean of 1,000 randomly picked subsets of 3 species from the genus and thus correct for the high number of species in some genera. Genera with less than 3 species were excluded from these analyses. Genome size estimates for a genus are based on the genome size of its species, which is defined as the majority set of functions of all strains for the functional universe measures. Pan genome openness is total pan genome size divided by mean genome size. Lines within violins show interquartile range and median.

### Supplementary tables captions

**Supplementary table 1. Selection of taxonomic analyses from the MGnify resource.** Taxonomic analysis are associated with runs, samples, and studies. Metadata of each datatype is shown. At most 1,000 samples were selected per annotated biome. For other selection criteria see **Methods**.

**Supplementary table 2. SNB for taxa across the tree of life and other features.** The database from which the feature is derived is indicated as the first word in square brackets, if no database is indicated the feature is derived from the MGnify data. Features with 'corrected' in the name refer to our completeness and contamination correction, see **Methods**. Genome size estimates and GC content features are calculated as the mean of all daughter species ('species' in the name) and taxonomy corrected (no 'species' in the name), see **Methods**. Pan genome features are calculated for all daughter species ('[all]' in the name) and for a random subset of 3 daughter species (1,000 random permutations per taxon) ('[3]' in the name).

**Supplementary table 3. Biases of genera towards high or low SNB.** Significant biases ($p < 0.05$) were decided based on the Mann-Whitney *U* test of the distribution of daughter genera versus the distribution of all other genera.

**Supplementary table 4. PATRIC genomes and associated data.** Features starting with 'genome.' are from the PATRIC database.

**Supplementary table 5. Gene Set Enrichment Analysis (GSEA) of genera sorted by SNB for different functional universes.** ES: enrichment score, NES: normalised enrichment score, FDR: false discovery rate. Tab 1: subsystem names, tab 2: subsystem subclasses, tab 3: GOs, tab 4: pathways.

**Supplementary table 6. Gene Set Enrichment Analysis (GSEA) of specialist genera (SNB < 0.35) sorted by alpha diversity for different functional universes.** ES: enrichment score, NES: normalised enrichment score, FDR: false discovery rate. Tab 1: subsystem names, tab 2: subsystem subclasses, tab 3: GOs, tab 4: pathways.
